# Cancer-associated fibroblasts promote drug resistance in *ALK*-driven lung adenocarcinoma cells by upregulating lipid biosynthesis

**DOI:** 10.1101/2023.08.08.552439

**Authors:** Ann-Kathrin Daum, Lisa Schlicker, Marc A. Schneider, Thomas Muley, Ursula Klingmüller, Almut Schulze, Michael Thomas, Petros Christopoulos, Holger Sültmann

## Abstract

Targeted therapy interventions using tyrosine kinase inhibitors (TKIs) provide encouraging treatment responses in *ALK*-rearranged lung adenocarcinomas, yet resistances occur almost inevitably. Apart from tumor cell-intrinsic resistance mechanisms, accumulating evidence supports a role of cancer-associated fibroblasts (CAFs) in affecting the therapeutic vulnerability of lung cancer cells. Here, we aimed to investigate underlying molecular networks shaping the therapeutic susceptibility of *ALK*-driven lung adenocarcinoma cells via tumor microenvironmental cues using three-dimensional (3D) spheroid co-culture settings. We show that CAFs promote therapy resistance of lung tumor cells against ALK inhibition by reducing apoptotic cell death and increasing cell proliferation. Using single-cell RNA-sequencing analysis, we show that genes involved in lipogenesis constitute the major transcriptional difference between TKI-treated homo- and heterotypic lung tumor spheroids. CAF-conditioned medium and CAF-secreted factors HGF and NRG1 were both able to promote resistance of 3D-cultured *ALK*-rearranged lung tumor cells via AKT signaling, which was accompanied by enhanced *de novo* lipogenesis and supression of lipid peroxidation. Notably, simultaneous targeting of ALK and SREBP-1 was able to overcome the established CAF-driven lipid metabolic-supportive niche of TKI-resistant lung tumor spheroids. Our findings highlight a crucial role of CAFs in mediating ALK-TKI resistance via lipid metabolic reprogramming and suggest new ways to overcome resistance towards molecular directed drugs by targeting vulnerabilities downstream of oncogenic signaling.

## Introduction

Lung adenocarcinomas, the most prevalent non-small cell lung cancer (NSCLC) subtype, can be stratified into different molecular subgroups based on recurrent genomic alterations. Among these, rearrangements of the *ALK* gene occur in approximately 6% of diagnosed cases (1). Crizotinib was the first approved tyrosine kinase inhibitor (TKI), which lead to remarkable responses in patients with advanced ALK+ NSCLC compared to standard chemotherapy (2, 3). Meanwhile, the next-generation ALK inhibitors (ALKi) brigatinib, alectinib and lorlatinib have become the preferred first-line treatment options based on superior systemic and intracranial efficacy compared to crizotinib in head-to-head randomized trials (4). However, the majority of patients who initally respond to these targeted drugs will show disease progression due to the emergence of drug resistance (5). This is caused by either genetic factors, i.e. secondary mutations in the ALK kinase domain or amplification of the ALK locus, with tumor cells being still ALK-dependent, or non-genetic factors, i.e., activation of bypass signaling pathways and phenotypic changes (6). While resistance mutations can be at least partly targeted by sequential treatment interventions (7), the mechanisms of resistance due to bypass signaling remain largely unknown.

Apart from tumor cell-intrinsic mechanisms, it has become evident that the tumor microenvironment (TME) also effectively shapes the therapeutic vulnerability of cancer cells (8, 9). Especially cancer-associated fibroblast (CAF) populations are known contributors to therapy resistance owing to a range of mechanisms including adhesion- and matrix-based signaling (10), paracrine crosstalk (11), immunosuppression (12) as well as metabolic reprogramming (13). The underlying signaling circuitry and connective tissue remodeling induces a disruption of normal tissue homeostasis and promotes cancer initiation and malignant progression. For example, paracrine interactions with stromal fibroblasts affect the sensitvity of NSCLC cells towards chemo- and targeted therapy (14, 15). However, the TME interactions leading to therapy resistance are not fully understood.

Metabolic reprogramming represents one of the hallmarks of cancer (16). Mounting evidence also supports an aberrant activation of the lipogenesis pathway in human cancers, which is required for membrane biogenesis and energy compensation of highly proliferative cancer cells (17–19). It is less clear, however, if this metabolic phenotype is driven exclusively by cancer cell-intrinsic processes or if it is also modulated by environmental factors, i.e. tumor-stromal interactions. The expression of genes encoding lipogenic enzymes is mainly controlled by the transcription factor family sterol regulatory element-binding proteins (SREBPs) (20–22), which occur in three different isoforms (SREBP-1a, SREPB-1c, SREBP-2). While SREBP-1 is primarily involved in regulating *de novo* lipogenesis, SREBP-2-driven gene activation affects cholesterol uptake and biosynthesis (22–24). *SREPB* expression and its transcription factor activity is tightly regulated by the cellular nutrient status in response to upstream signaling networks (e.g. PI3K/AKT/mTORC1 pathway) (25–28). Likewise, receptor tyrosine kinase (RTK) signaling gives rise to rewiring of lipid metabolism in oncogene-driven cancers (29–32). In line with this, Chen et al. (33) demonstrated perturbations of lipid metabolism following TKI-treatment in EGFR-driven NSCLCs. The same study also found that sustained SREBP-1-dependent lipogenesis was a key mediator of resistance to EGFR-targeted therapy, which could be reversed by metabolic interventions using SREBP-1 inhibition.

Since to our knowledge no prior studies have investigated tumor stroma-dependent global gene expression changes in TKI-treated EML4-ALK-positive NSCLC, the aim of our study was to explore tumor-stroma-related processes leading to non-genetic resistance in this tumor type. In order to mimic fibroblast-tumor cell interactions in the TME *in vitro,* we established spheroid co-culture models of ALK-driven NSCLC cell lines and CAFs. We employed single-cell RNA-sequencing (scRNA-seq) to analyze differential gene expression between TKI-treated homo- and heterotypic lung tumor spheroids. Our analyses lead us to focus on the deregulation of tumor cell lipid metabolism evoked by factors in the TME but also provide a resource of data for further characterization of non-genetic TKI therapy resistance in these models.

## Results

### The presence of CAFs reduces apoptosis and promotes cell cycle progression of lung tumor cells upon ALK inhibition

CAFs are known promoters of tumor cell resistance against systemic chemotherapy and targeted therapy interventions (8). In order to investigate the stroma-mediated drug sensitivity of NSCLC tumor cells we established 3D cultures of *EML4-ALK* translocation carrying H2228 (E6;A20) as well as H3122 (E13;A20) cells with and without CAFs (Fig. 1A). The CAF phenotype was induced via stimulation with 2 ng/ml TGF-β1 using MRC-5 fibroblasts and two primary fibroblast lines (FB1, FB2) prior to co-cultivation. CAF activation was confirmed on a phenotypic and molecular level by a stellate cell-shaped morphology (Supplementary Fig. S1A) as well as elevated expression levels of α-smooth muscle actin (αSMA) and fibroblast activation protein (FAP) (Supplementary Fig. S1B and C), respectively. Treatment with appropriate concentrations (IC_75_ values, Supplementary Table S1) of brigatinib and lorlatinib, as obtained by dose-response modeling (Supplementary Fig. S2), led to an increase of dead and apoptotic cells in H2228 (Fig. 1B, Supplementary Fig. S3A and B) and H3122 (Fig. 1C, Supplementary Fig. S3C) spheroids as assessed by Annexin V+/FVD staining and subsequent flow cytometric analysis. In contrast, direct co-cultivation with activated MRC-5, FB1, or FB2 cells, respectively, significantly reduced tumor cell death upon TKI application when compared to spheroids consisting of tumor cells only (Fig. 1B and C). Of note, the tumoricidal effect of both drugs appeared to be stronger in in case of the EML4-ALK variant 1 (v1)-driven H3122 cells compared to the EML4-ALK v3-driven H2228 (Fig. 1B and C, Supplementary Fig. S2B and C). Next, the combinatorial staining of the proliferation associated antigen Ki-67 and propidium iodide (PI) revealed significant changes in the cell cycle of homotypic H2228 (Fig. 1D, Supplementary Fig. S4A and B) and H3122 (Fig. 1E, Supplementary Fig. S4C) spheroids and a switch towards a G0 arrest following brigatinib or lorlatinib treatment, respectively. In contrast, exposure of heterotypic H2228 and H3122 spheroids to both ALKi’s, lead to significantly increased cell cycle progression of the tumor cells in a CAF-dependent manner. Together, these results provide strong evidence that CAFs establish a protective niche for *EML4-ALK*-rearranged lung cancer cells towards ALK-inhibition.

**Figure 1:**
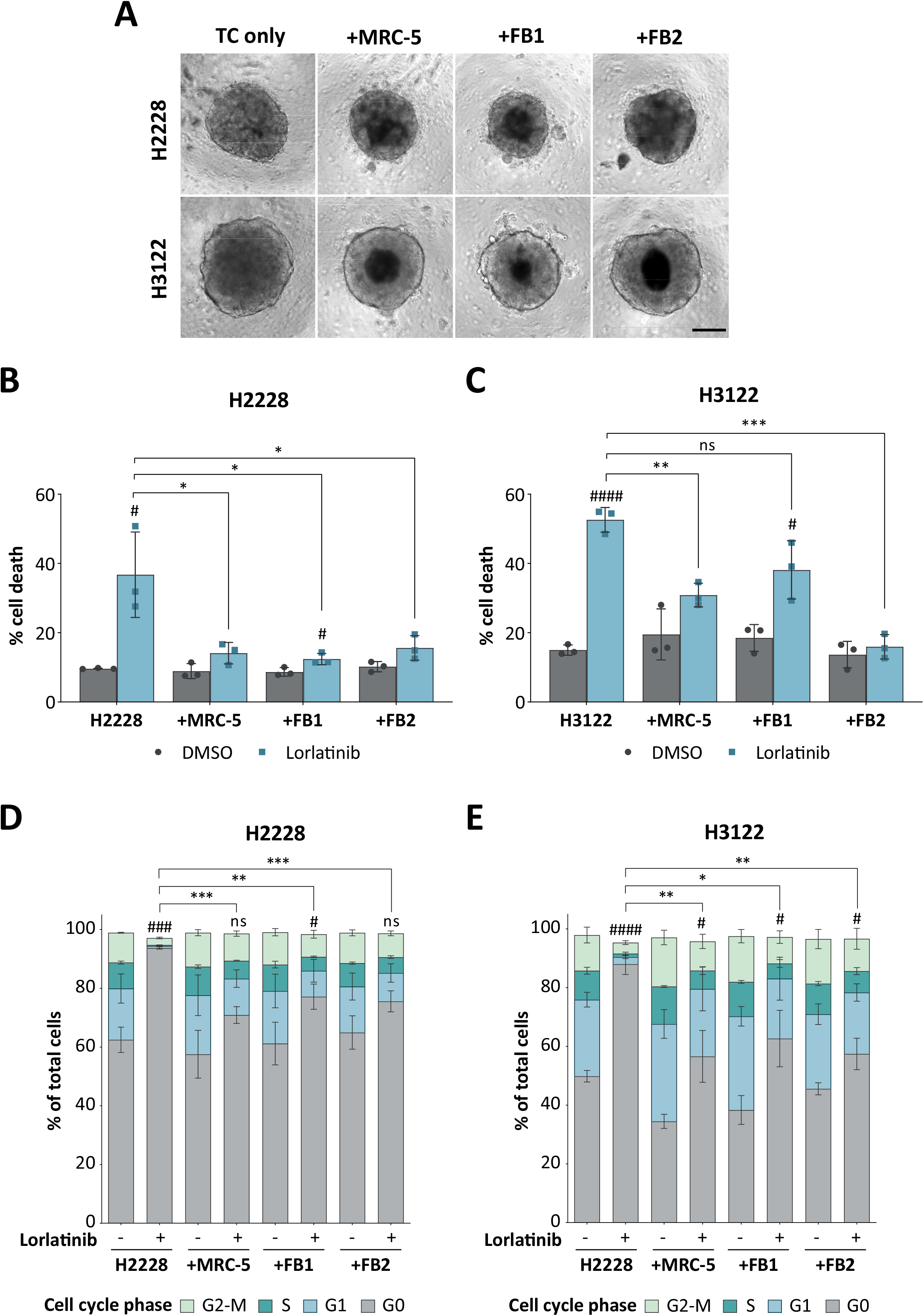
(A) H2228 or H3122 lung tumor cells (TC) were seeded alone or in combination with three TGF-β1-activated fibroblast lines (MRC-5, FB1, FB2) for the generation of homo- and heterotypic lung tumor spheroids, respectively. Scale bar: 200 µm. Quantification of cell death rates of H2228 (B) and H3122 (C) cells as given by the sum of early and late apoptotic/necrotic cells following flow cytometric analysis (n = 3). Data are presented as mean ± SD. #,p≤0.05; ###, p≤0.001 compared to corresponding DMSO controls. ns, not significant; *, p≤0.05; **, p≤0.01 in comparison to lorlatinib-treated mono-cultures. FVD, fixable viability dye. Quantification of the portion of H2228 (D) and H3122 (E) cells according to their cell cycle phase status (n = 3). Data are presented as mean ± SD. ns, not significant; #,p≤0.05; ###, p≤0.001; ####, p≤0.0001 of cells wihtin the G0-phase compared to corresponding DMSO controls. *, p≤0.05; **, p≤0.01; ***, p≤0.001 of cells wihtin the G0-phase in comparison to lorlatinib-treated mono-cultures. PI, propidium iodide.

### Distinct clusters of ALK+ NSCLC cells and CAFs as revealed by scRNA-seq

To analyze the molecular processes in tumor cells and CAFs, we applied scRNA-seq of dissociated H2228 and H3122 tumor spheroids either mono-cultured or co-cultured with TGF-β1-activated FB2 fibroblasts. We acquired transcriptome data for a total of 86,636 individual cells, with an average of 4,600 unique genes detected per cell (Supplementary Table S2). To determine differentially expressed genes, we employed a series of analytical steps for each individual dataset including quality control and filtering, principal component analysis (PCA) and a graph-based clustering approach. Dimensionality reduction and visualization of cell-type clusters defined by their transcriptional profiles were performed by using uniform manifold approximation and projection (UMAP). Seven clusters (numbered 0 to 6) were observed for both lorlatinib-treated H2228 (Fig. 2A) and H3122 (Fig. 2B) spheroid-derived samples. To characterize inter- and intra-population heterogeneity and define the identity of the clusters, pairwise differential gene expression analysis for each cluster against all others was performed to generate cluster-specific marker genes conserved across all conditions. A total of 1,161 (H2228) and 1,000 (H3122) genes were identified that best grouped the cells into the seven subgroups. Heatmaps illustrating the ten most differentially expressed genes of each cell cluster are shown in Supplementary Fig. S5. Known markers associated with matrix remodeling (e.g. *TIMP1, COL1A2*), fibroblast activation (e.g. *IGFBP5, FAP*) as well as fibroblast-derived growth factors (e.g. *HGF, FGF2*) and cytokines (e.g. *CCL2*) clearly identified the transcriptionally distinct cluster 2 (H2228, Supplementary Fig. S5A) or cluster 3 (H3122, Supplementary Fig. S5B) cells as the CAF population. This was supported by the fact that these clusters were present only in the co-culture samples (Fig. 2A and B). The H2228 and H3122 tumor cell fractions were divided into six sub-clusters based on UMAP analysis. This heterogeneity could be partly explained by the presence of cycling and proliferating cells in cluster 3 and 4 within the H2228 sample group (Supplementary Fig. S5A) as well as cluster 1 and 4 of H3122 samples (Supplementary Fig. S5B). This was further supported by the specific expression of cell proliferation (e.g. *TOP2A, PCLAF, CENPF*) and cell cycle (e.g. *PTTG1, CCNB1*) markers. The abundance of cells within these cycling clusters also exhibited a clear decrease in lorlatinib-treated mono-culture-derived samples compared to all other samples, which corroborates the findings of the previous cell cycle analysis (see Fig. 1D and E).

**Figure 2:**
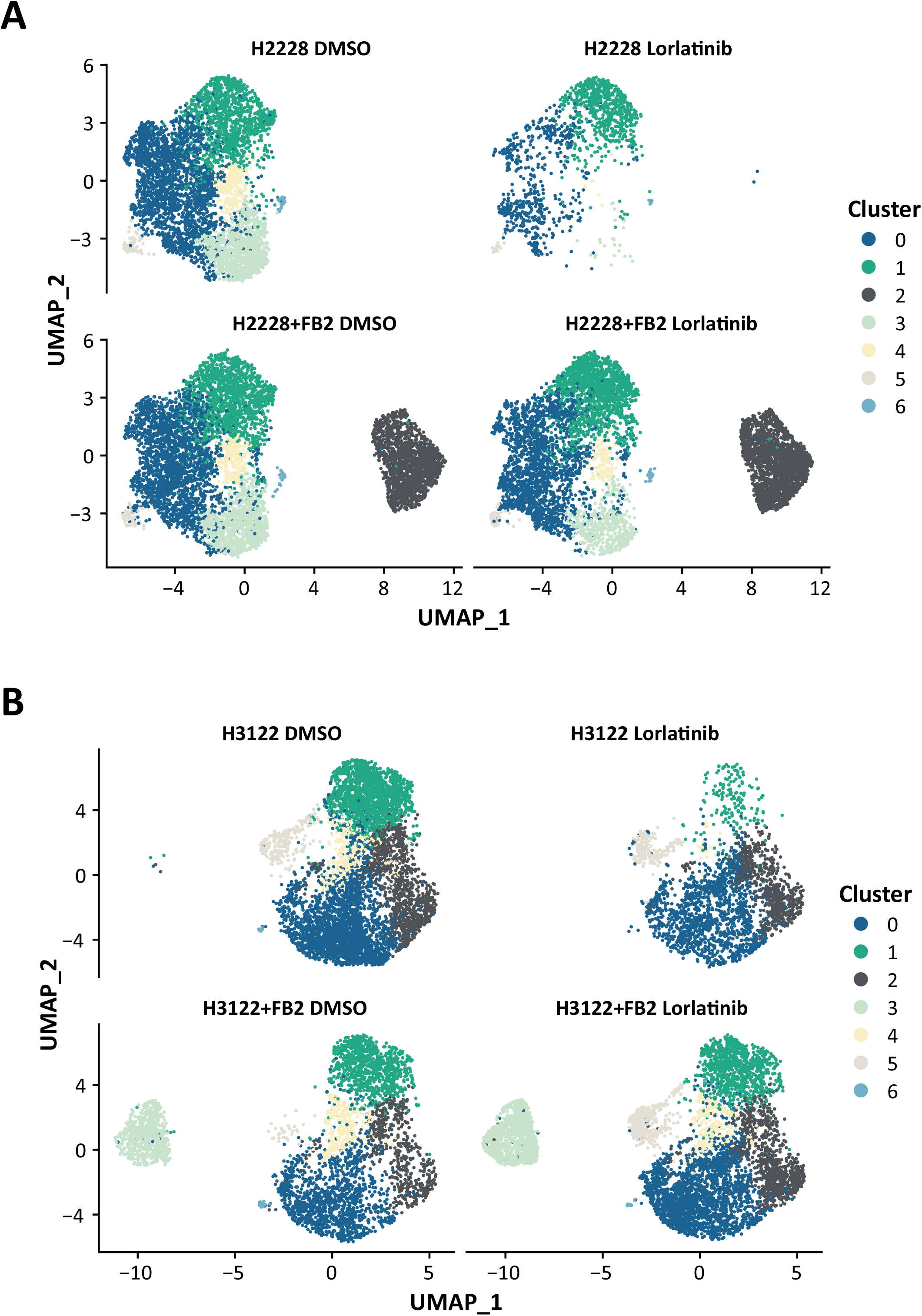
Uniform manifold approximation and projection (UMAP) representation of lorlatinib-treated H2228 (A) and H3122 (B) cells following graph-based clustering colored by cluster identity.

### Co-cultivation with CAFs drives upregulation of the lipid metabolic pathway in ALKi-treated lung tumor spheroids

In order to explore the molecular processes facilitating survival of ALK-driven lung cancer cells under the influence of CAFs, differential gene expression analysis between treated mono- and co-culture conditions was performed. After exclusion of CAF clusters, 334 to 609 differentially expressed genes (avg. logFC |≥ 0.25|) were identified in brigatinib or lorlatinib-treated H2228 and H3122 mono-cultures compared to co-culture conditions, respectively (Supplementary Fig. S6A). Of these, considerable overlaps of upregulated genes in TKI-treated lung tumor cells in response to CAF co-cultivation compared to mono-cultures were observed (Supplementary Fig. S6B and S7). These differentially expressed genes were subjected to Gene Set Enrichment Analysis (GSEA) based on the Molecular Signatures Database (MSigDB) which revealed enriched pathways related to cell proliferation, metabolic activity and signal transduction based on Rho-GTPases (Fig. 3A). Moreover, among the enriched biological processes and pathways related to metabolic activities, the most striking terms were linked to lipid metabolism. A selection of genes encoding for lipogenic regulators are highlighted in Fig. 3B and Supplementary Fig. S8A, demonstrating upregulation of *de novo* lipogenic enzymes in TKI-treated co-cultures when compared to corresponding mono-culture conditions. Of note, the gene coding for stearoyl-CoA desaturase-1 (*SCD*) a key regulatory factor of fatty acid (FA) synthesis, was among the top 79 upregulated genes. *SCD* is a well-established transcriptional target of sterol regulatory binding protein 1 (SREBP-1), encoded by the sterol regulatory element-binding transcription factor 1 (*SREBF1*) gene, whose downregulation following ALK-TKI treatment was also prevented by co-culture with CAFs. The differential expression of ATP citrate lyase (*ACLY*), acetyl-CoA carboxylase alpha (*ACACA*), fatty acid synthase (*FASN*), *SCD,* and *SREBF1* was further validated by RT-qPCR subsequent to magnetic-based cell sorting of co-cultured H2228 and H3122 tumor spheroids (Fig. 3C, Supplementary Fig. S8B). These results indicate that upregulation of fatty acid metabolism might contribute to CAF-dependent resistance towards ALK inhibition.

**Figure 3:**
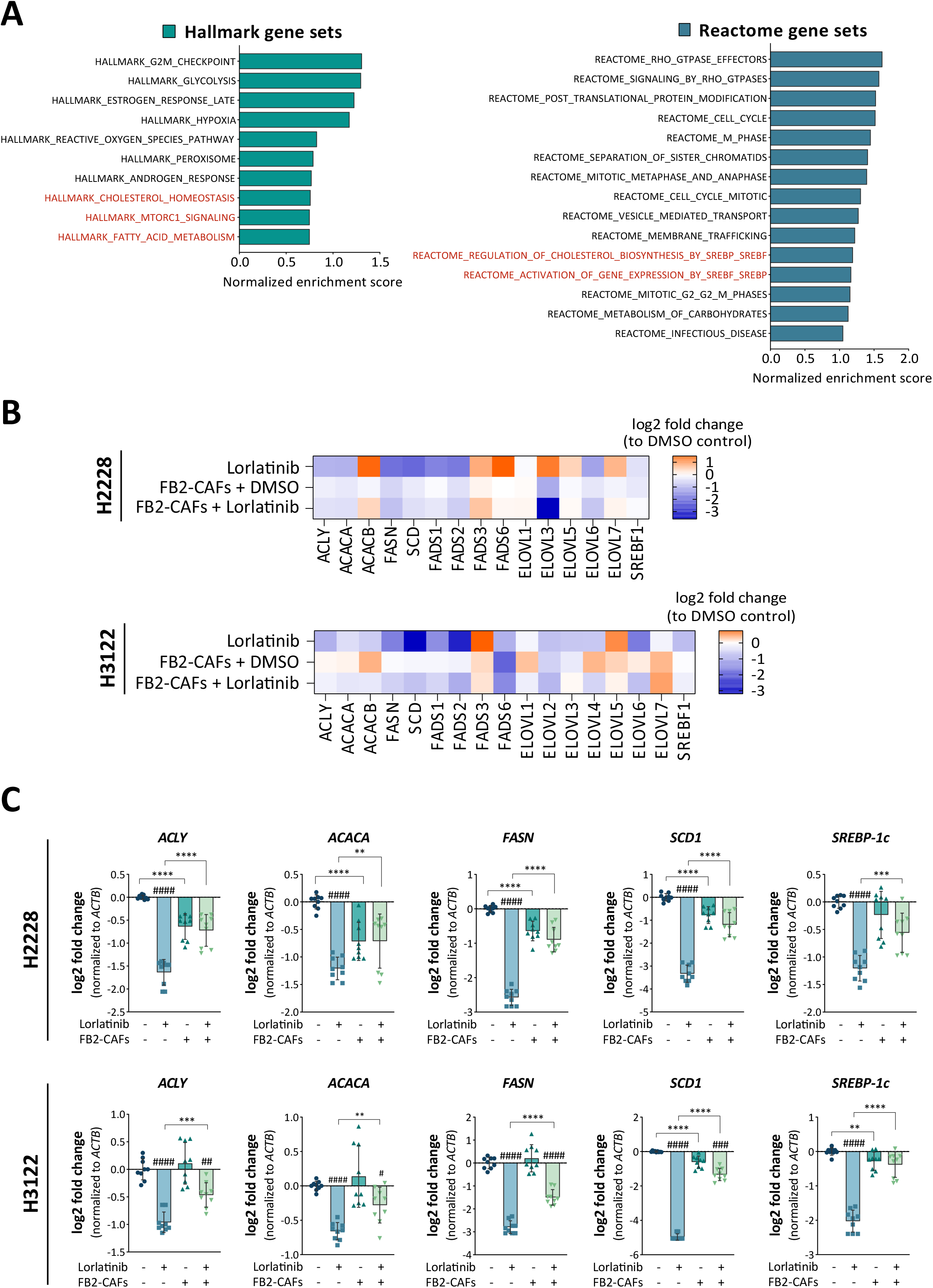
(A) Top deregulated pathways of hallmark (green bars) and reactome (blue bars) gene sets derived from the Molecular Signatures Database (MSigDB) in FB2 co-cultured versus mono-cultured H2228 and H3122 cells following ALK inhibition based on gene set enrichment analysis (GSEA) (FDR ≤ 0.25). (B) Expression heatmap of fatty acid metabolism-related genes in single-cell transcriptome datasets of lorlatinib-treated H2228 and H3122 cells. (C) Co-cultivation with FB2-CAFs influences expression of fatty acid metabolism-related targets upon ALK signaling perturbation using lorlatinib in H2228 and H3122 cells (n = 3). Data are presented as mean ± SD. #, p≤0.05; ##, p≤0.01; ###, p≤0.001; ####, p≤0.0001 compared to corresponding DMSO controls. **, p≤0.01; ***, p≤0.001; ****, p≤0.0001 in comparison to lorlatinib-treated mono-cultures.

### Ligand-receptor interaction analysis predicts CAF-associated ligands that convey therapy resistance in ALK+ lung cancer cells

To identify secreted factors and their cognate receptors for paracrine-mediated signaling, an *in silico* analysis of the scRNA-seq data was conducted by employing the ‘RNA-magnet’ algorithm (34). Putative tumor-stroma interactions were inferred based on a curated list of ligand-receptor pairs and the enrichment of ligand:receptor transcripts between the CAF population and each tumor cell cluster. The results of the top predicted interactions pairs (interaction score > 0.2) across all conditions of both H2228 and H3122 samples are listed in Supplementary Table S3. Among the computed interactions, key paracrine ligands derived from CAFs comprised RTK-ligands such as HGF, FGF2 and NRG1 as well as Wnt-signaling associated factors (e.g. WNT5A, SFRP1). Of these, the highly abundant HGF was predicted to bind to its cognate receptor MET, but also to SDC1 and ST14 (Supplementary Table S3, Fig. 4). HGF is a widely known CAF-secreted factor and the HGF/MET-axis has often been found to be associated with cancer progression and the induction of therapy resistance (35, 36). Secreted WNT5A was suggested to interact with a wide range of tumor cell-expressing receptors, such as LRP5, LRP6, FZD5, and FZD6 (Supplementary Table S3, Fig. 4). WNT5A, an activator of non-canonical Wnt pathways, exhibits pro-tumorigenic functions and contributes to inflammation and immunosuppression in the tumor microenvironment (37). Another interesting computed stromal-epithelial interaction pair was NRG1-ERBB3 (Supplementary Table S3, Fig. 4). The CAF-derived factor NRG1 was demonstrated to modulate therapy response in a variety of cancers (38, 39). Of note, NRG1 signaling via ErbB receptors is involved in the (lipid-demanding) myelination by Schwann cells (40, 41) and oligodendrocytes (42). In addition, interactions of the CAF-derived decoy receptor TNFRSF11B/OPG with the apoptosis-inducing ligand TNFSF10/TRAIL were suggested. Interestingly, OPG was previously reported to protect cancer cells from TRAIL-induced apoptosis (43, 44).

**Figure 4:**
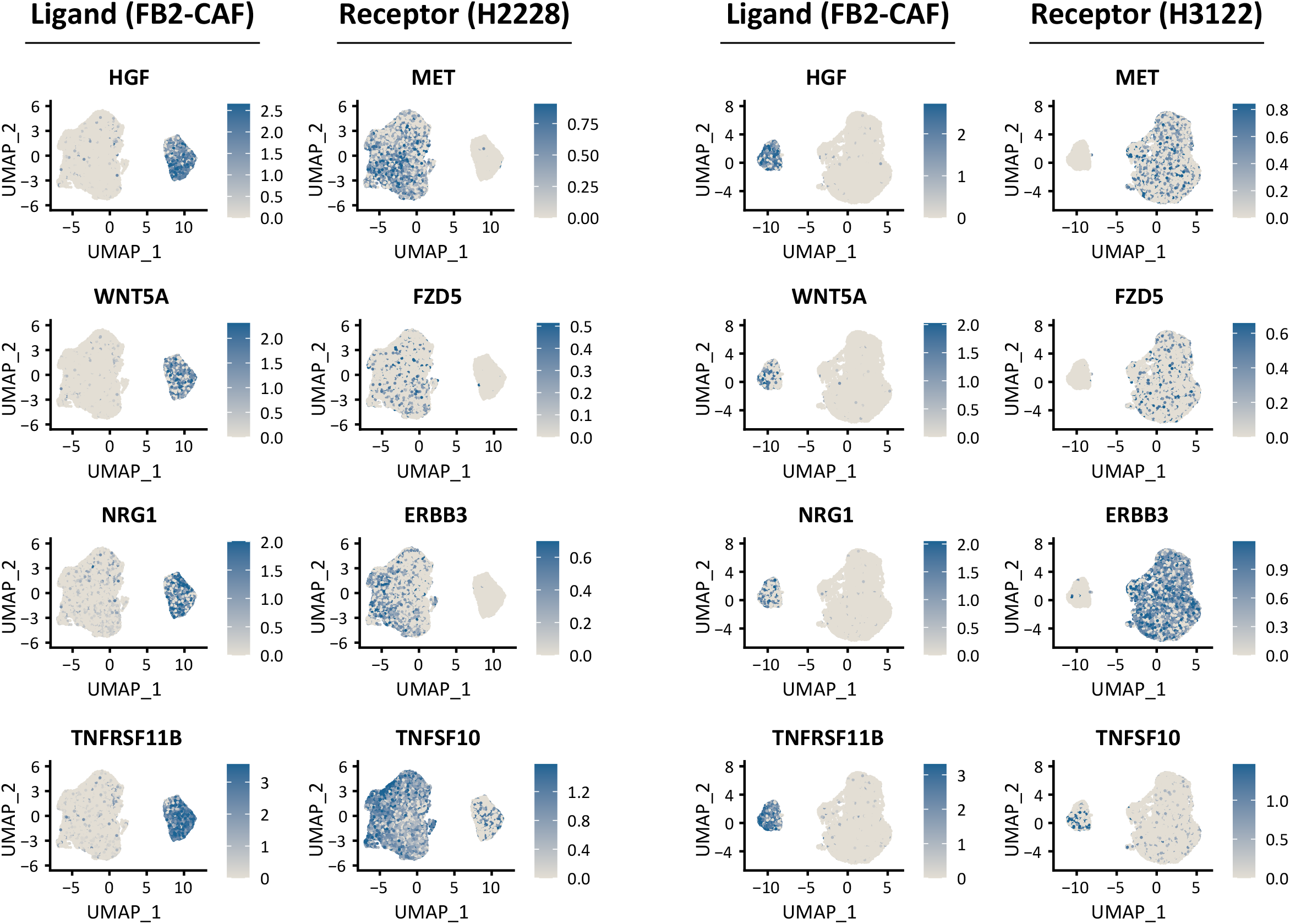
Inference of cellular interactions from single-cell transcriptome data using ‘RNA-Magnet’. Routes of cell-cell communication between FB2-CAFs and H2228 or H3122 lung tumor cells were predicted following ligand-receptor interaction analysis. Selected ligand-receptor pairs are visualized as feature plots.

The ligand-receptor interaction analysis identified a variety of CAF-secreted factors, which could act as potential communicators between CAFs and lung tumor cells and thus as initiators for stromal-driven therapy resistance. To corroborate this, spheroid growth analyses using combinatorial treatment experiments with ALK-TKIs and CAF-conditioned medium (CM) derived from TGF-β1-activated and 3D-cultured lung FB2 fibroblasts demonstrated increased growth of H2228 (Supplementary Fig. S9A and B) and H3122 (Supplementary Fig. S9A and C) spheroids as opposed to treatment with brigatinib or lorlatinib alone. These experiments indicated that the CAF supernatants contain molecular factors promoting treatment resistance of the tumor cells.

To validate the expression of selected CAF-associated ligands inferred via interaction analysis, the active secretion of HGF, NRG1β1, TNFRSF11B/OPG, and WNT5A in cell culture supernatants derived from previously 3D-cultured MRC-5, FB1 and FB2 fibroblasts was measured using multi-analyte profiling. All tested ligands were detected in conditioned medium obtained from both untreated and TGF-β1-treated fibroblasts (Fig. 5A). Here, differences in analyte concentrations were observed in a fibroblast line- and activation status-dependent manner. Of note, the *NRG1*-gene gives rise to numerous protein isoforms (e.g. NRG1α and NRG1β1) through alternative promoter usage and splicing (45). The total NRG1 concentration is therefore likely higher in the fibroblast CM than the detected 0.02-0.7 ng/ml of NRG1β1 only. Interestingly, the secretion level of all analytes tested was increased following TGF-β1-mediated activation in MRC-5, FB1 and FB2 fibroblasts. In contrast, activated MRC-5 fibroblast secreted less HGF compared to their native counterpart, a finding which is consistent with previous studies (46, 47).

**Figure 5:**
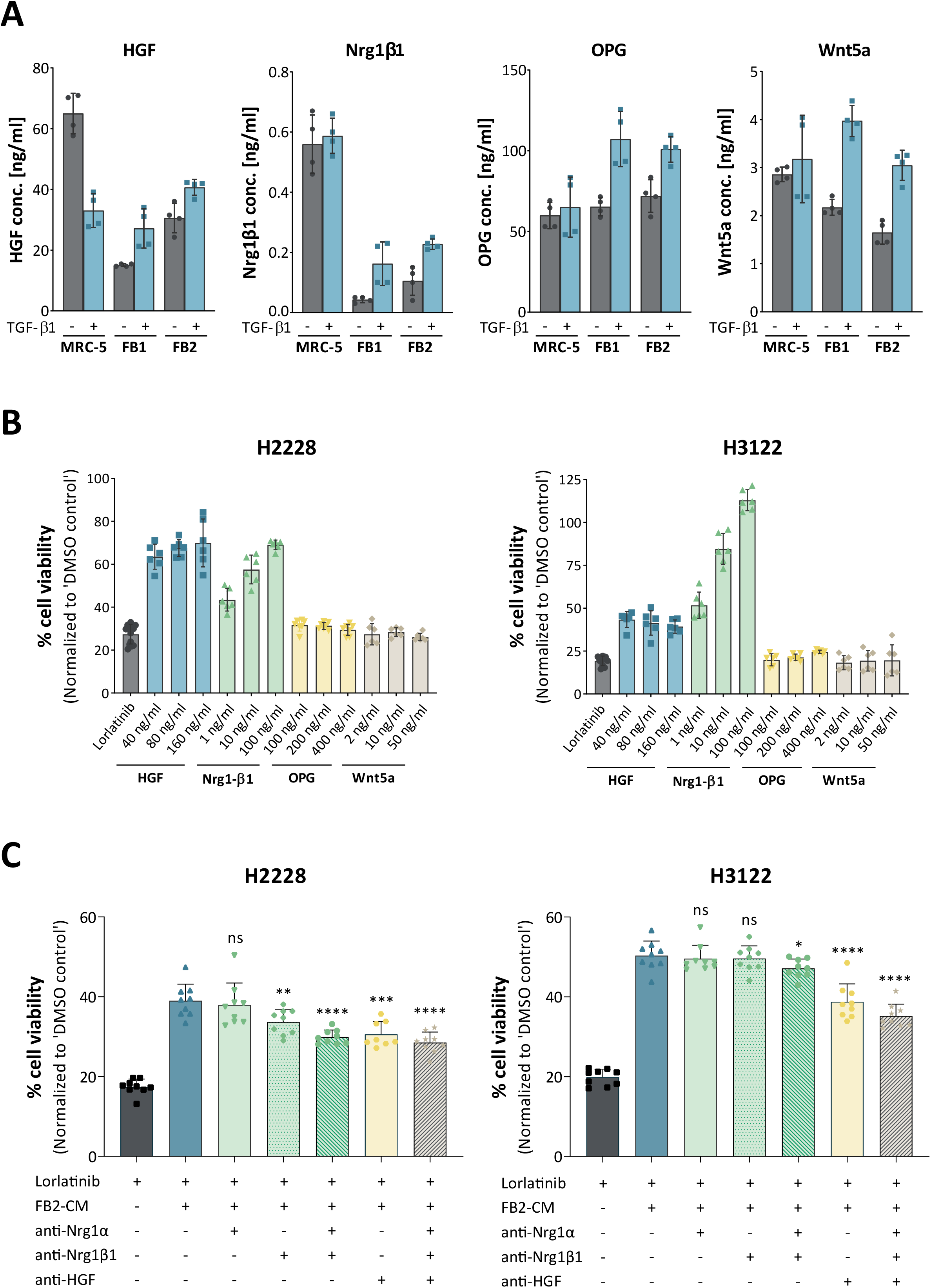
(A) Validation of CAF-secreted candidate ligands using Luminex and ELISA. Concentrations were measured according to generated standard curves and are given for native versus TGF-β1-activated fibroblasts (n = 2). (B) Cell viability analysis of ALK-inhibited lung tumor cells following treatment with CAF-associated ligands (n = 2). (C) Assessment of cell viability of lorlatinib-treated lung tumor spheroids subsequent to addition of FB2-conditioned medium (CM) and inhibitory antibodies directed against the CAF-associated ligand Nrg1α, Nrg1-β1, and HGF (n = 3). All data are presented as mean ± SD. ns, not significant; *, p≤0.05; **, p≤0.01; ***, p≤0.001; ****, p≤0.0001 in comparison to treatment with lorlatinib and FB2-CM. HGF, hepatocyte growth factor; NRG1, neuregulin 1; TGF-β1, transforming growth factor-beta 1; OPG, osteoprotegerin.

To address the possibility, that the selected ligands alone could have mechanistic consequences on lung tumor cell drug sensitivity, recombinant HGF, NRG1β1, OPG, and WNT5A proteins were applied to ALKi-treated H2228 and H3122 tumor spheroids. Lorlatinib treatment considerably reduced cell viability compared to the vehicle control (Fig. 5B). OPG and WNT5A appeared to possess no paracrine potency in mediating tumor cell drug sensitivities in all tested conditions. Since these ligands were secreted by all fibroblast lines, a pivotal role for pro-fibrotic autocrine signaling purposes cannot be excluded. Addition of HGF or NRG1β1, however, could partially or fully reverse the TKI-induced decrease in H2228 and H3122 cell viability. This effect was observed in a concentration-dependent manner for NRG1β1, whereas increasing HGF concentrations did not result in a cumulative tolerability of H2228 and H3122 spheroids towards ALK inhibition. In line with this, the resistance-mediating effect of FB2-CM addition to lorlatinib-treated H2228 and H3122 tumor spheroids was partially inhibited by neutralizing antibodies directed against HGF, NRG1α, and NRG1β1 as well as combinations thereof (Fig. 5C). Taken together, these data established a direct correlation of CAF-associated paracrine factors in conferring therapy resistance to H2228 and H3122 cancer cells.

### The CAF-derived secretome upregulates lipid metabolism and AKT signaling in ALKi-treated lung tumor spheroids

The differential expression of lipogenic enzymes upon ALK inhibition could be validated in direct co-culture conditions. Since the CAF-derived secretome was also able to interfere with the drug susceptibilty of ALK-driven lung tumor spheroids, we hypothesized that a potential link existed between CAF-mediated paracrine signaling and an enhanced lipid metabolic activity in the tumor cells. Western blot analysis of selected lipogenic enzymes suggested, that Lorlatinib caused a decrease in ACLY and FASN concomitant with reduced expression of their upstream transcription factor SREBP-1. The addition of CAF-CM (Fig. 6A) as well as of the single CAF-associated ligands HGF and NRG1β1 (Fig. 6B) was able to partially restore the observed protein expression changes, respectively.

**Figure 6:**
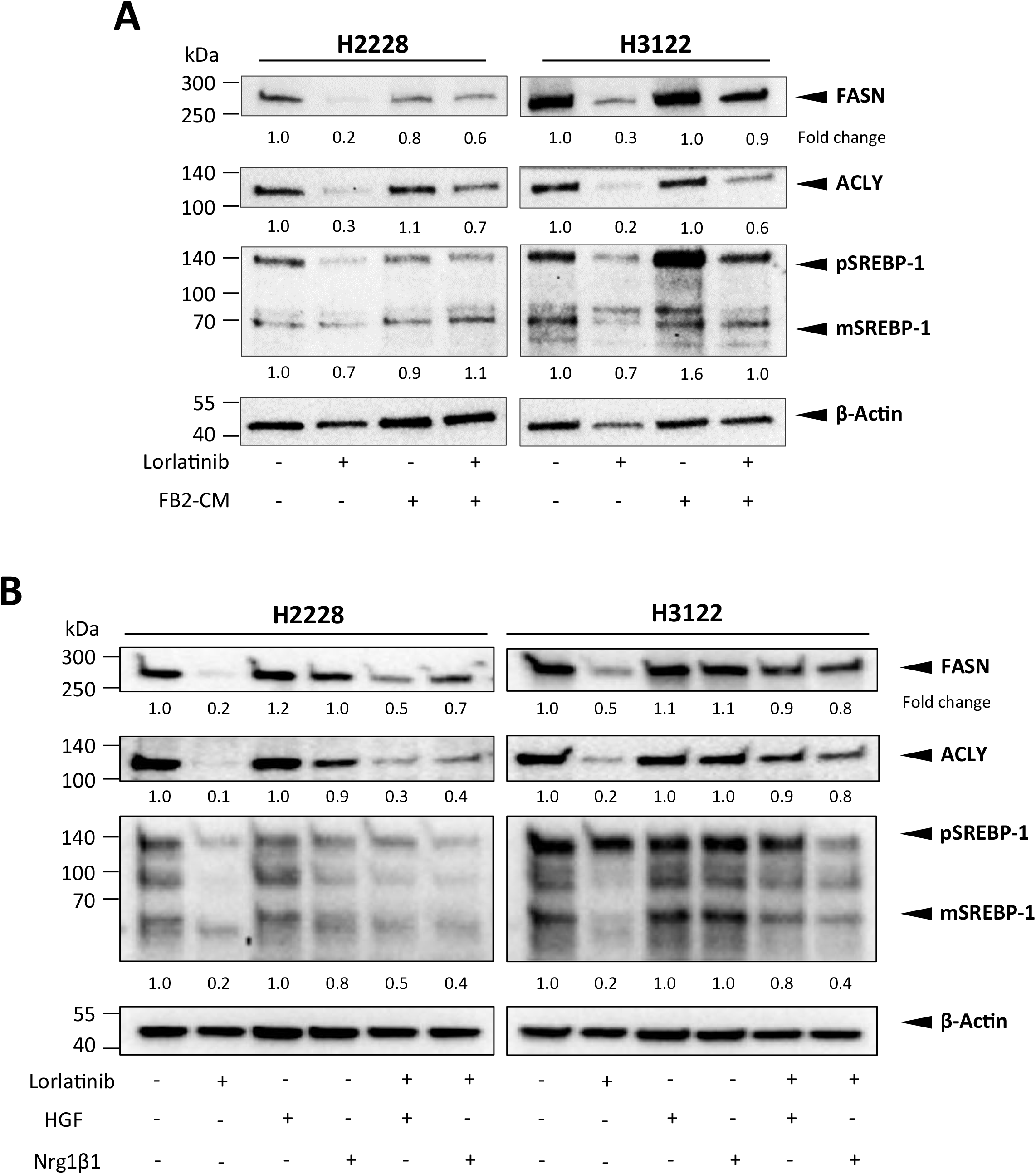
FB2-conditioned medium (CM) (A) and CAF-associated ligands HGF and Nrg1-β1 (B) influence expression of fatty acid metabolism-regulating targets upon ALK signaling perturbation using lorlatinib in H2228 and H3122 cells as shown by representative western blots of FASN, ACLY, and SREBP-1 in H2228 and H3122 cells. Expression fold changes were normalized to β-Actin. ACLY, ATP-citrate lyase; FASN, fatty acid synthase; pSREBP-1, precursor sterol-responsive element binding protein-1; mSREBP-1, mature SREBP-1.

The PI3K-AKT pathway is a commonly activated signaling pathway downstream of mutant ALK (48). Previous work demonstrated an induction of lipid biosynthesis via PI3K-AKT-mTOR signaling, with mTORC1 being a direct regulator of SREBP transcription factor activity (49, 50). We therefore investigated whether the fibroblast-derived secretome enhances AKT signaling in ALK-inhibited lung tumor spheroids. While ALK inhibitor treatment lead to a reduction of phosphorylated AKT and the mTOR downstream target S6K1, AKT signaling was effectively re-induced following cultivation of H2228 and H3122 tumor spheroids with CAF-CM (Fig. 7A).Treatment with exogenous HGF or NRG1β1 was also sufficient to induce phosphorylation of AKT and S6K1 despite inhibition of oncogenic ALK (Fig. 7B). These data imply that the CAF-derived secretome confers therapy resistance via induction of the PI3K-AKT-mTOR signaling axis and thereby drives the expression of SREBP-targets.

**Figure 7:**
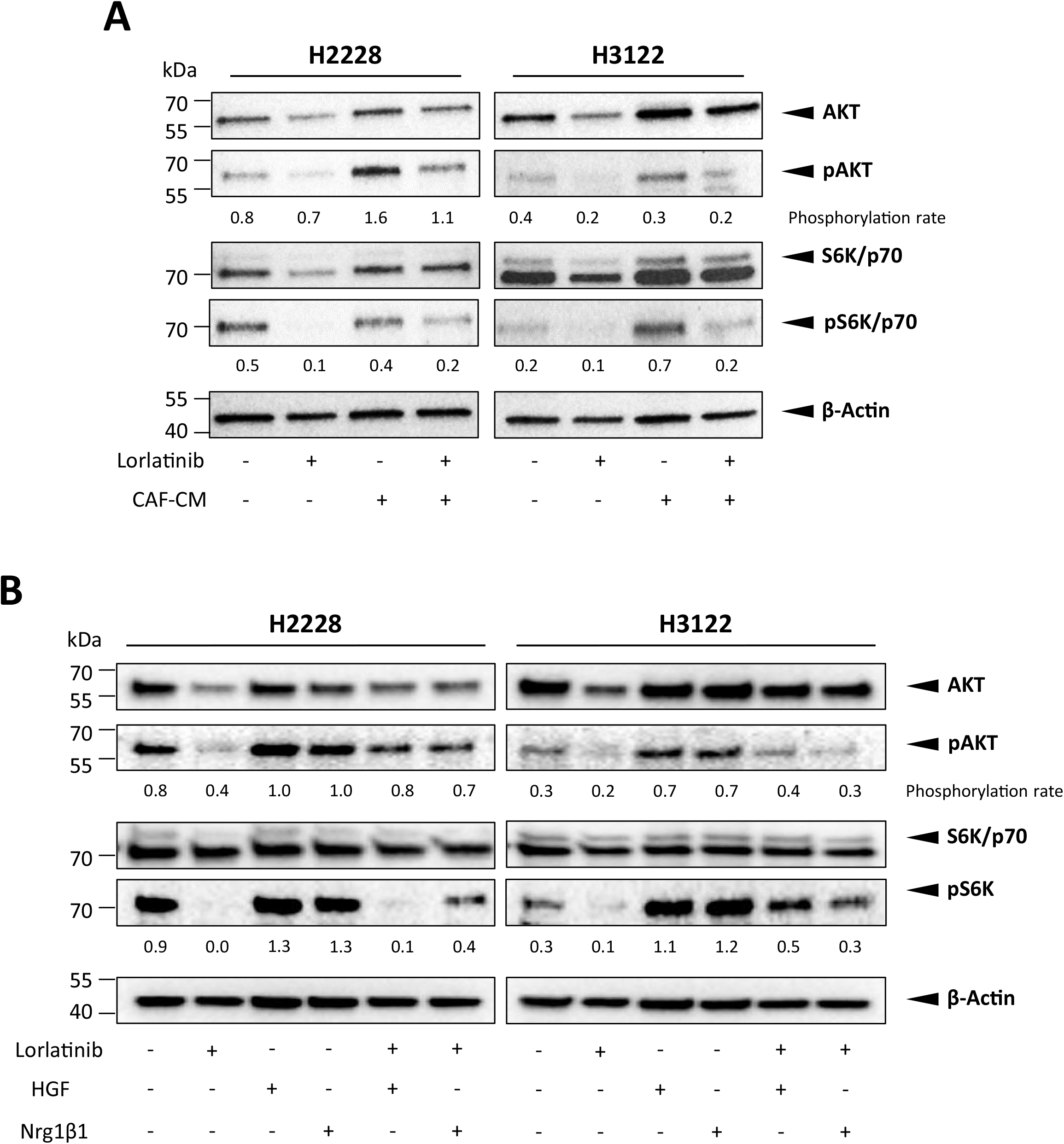
FB2-conditioned medium (CM) (A) and CAF-associated ligands HGF and Nrg1-β1 (B) influence acitvation of AKT signaling upon ALK signaling perturbation using lorlatinib in H2228 and H3122 cells as shown by representative western blots of AKT/pAKT and S6K/pS6K in H2228 and H3122 cells. Phosphorylation rates were normalized to β-Actin. AKT, protein kinase B; S6K, ribosomal S6 kinase.

### CAF-CM rescues lung tumor spheroids from TKI-induced interference with de novo lipogenesis and lipid peroxidation

Since ALK inhibition evokes differential expression of lipid metabolizing enzymes, we performed mass spectrometry-based phospholipidome profiling to evaluate changes in the proportion of lipid species. Inhibition of oncogenic ALK lead to an increase of poly-unsaturated fatty acids (PUFAs) of the phosphatidylethanolamine (PE, Fig. 8A) and phosphatidylcholine (PC, Fig. 8B) phospholipid species at the expense of saturated and mono-unsaturated phospholipids, a typical observation following inhibition of *de novo* lipogenesis (51). Addition of CAF-CM to H3122 lung tumor spheroids was able to partly abrogate this shift towards higher levels of poly-unsaturated lipid.

**Figure 8:**
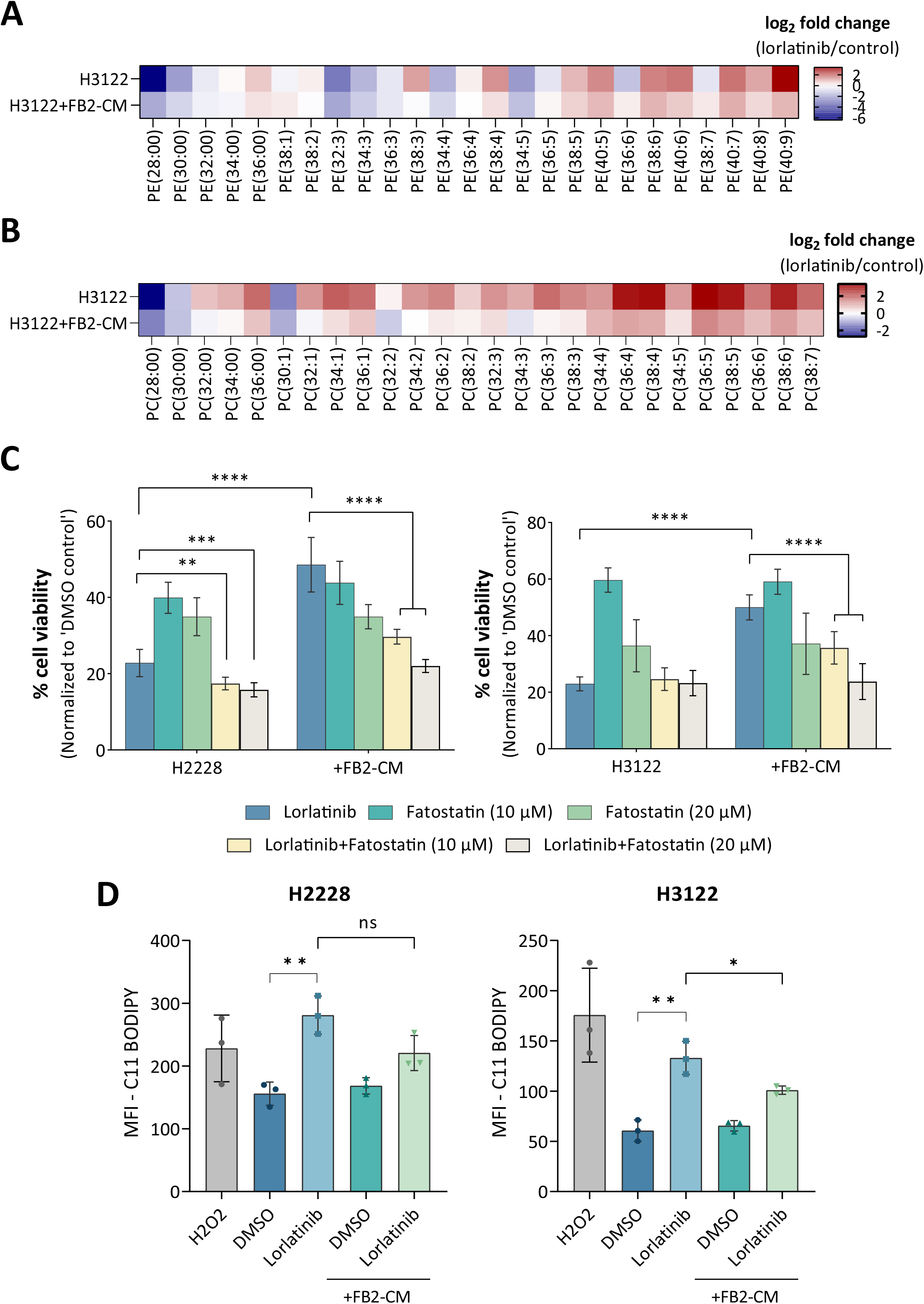
(A) Heatmap of log_2_ ratios of phosphatidylethanolamine (PE)-species in lorlatinib-treated H3122 cells w/ and w/o FB2-conditioned medium over vehicle treated cells. PE-species are indicated by their total number of fatty acid carbons, followed by the total number of unsaturations (n = 3). (B) Heatmap of log_2_ ratios of phosphatidylcholine (PC)-species in lorlatinib-treated H3122 cell w/ and w/o FB2-conditioned medium over vehicle treated cells. PC-species are indicated by their total number of fatty acid carbons, followed by the total number of unsaturations (n = 3). (C) Cell viability analysis following combinatorial treatment of H2228 and H3122 tumor spheroids with fatostatin and lorlatinib (n = 3). (D) Quantification of lipid peroxidation using C11 BODIPY. Hydrogen peroxide (H_2_O_2_) served as positive control. All data are presented as mean ± SD. ns, not significant; *, p≤0.05; **, p≤0.01; ***, p≤0.001; ****, p≤0.0001.

To further determine the importance of lipogenesis on the cell proliferative response of TKI-resistant ALK+ lung cancer cells, we evaluated the effect of fatostatin, a small molecule inhibitor which hinders trafficking of SREBP-1 to the Golgi compartment, and thereby its proteolytic cleavage and activation. Viability assays showed that SREBP-1 inhibition through fatostatin sensitized lung tumor spheroids to ALK inhibition despite the presence of CAF-CM (Fig. 8C).

The above findings suggest that CAF-driven therapy resistance is a result of enhanced lipid saturation and thus decreased lipid peroxidation, since PUFAs are more prone to lipid peroxidation than saturated phospholipids (51). To test this hypothesis, we measured lipid reactive oxygen species (ROS) accumulation using BODIPY 581/591 C11 fluorescent staining. Flow cytometry analysis showed a strong increase in lipid peroxidation subsequent to lorlatinib treatment in both, H2228 and H3122 spheroids (Fig. 8D). In contrast, tumor spheroids treated with CAF-CM accumulated less lipid peroxides despite inhibition of ALK.

Collectively, these data show that interference with ALK signaling inhibits *de novo* lipogenesis concomitant with an increase in lipid peroxidation, an effect which is attenuated in the presence of CAF-CM. In addition, SREBP-1 inhibition sensitizes therapy-resistant lung cancer cells to ALK-targeting therapy.

## Discussion

Altough precision oncology has made significant strides in tailoring therapy to the molecular alterations of individual tumors, acquired resistance to targeted drugs remains a key clinical challenge (3). One opportunity for cancer cells to bypass drug response is through cellular crosstalk with frequently abundant CAF populations. Thus, exploiting vulnerabilities occuring in the presence of CAFs represents an attractive strategy to overcome therapy resistance. In order to enhance the pathophysiological relevance of *in vitro* modeling, the co-culture model devised in this study involved cultivation of EML4-ALK-driven lung adenocarcinoma cell lines and CAFs in a 3D setting. Generated lung tumor spheroids were utilized to explore signaling networks between the two heterologous cell types shaping the susceptibility to molecular-directed drugs via microenvironmental cues. In kinase-addicted NSCLC, a reduction of tumor cell survival following EGFR- and ALK-directed therapy interventions could be counteracted with exposure to non-malignant CAF populations or CAF-associated paracrine mediators (52–55). This well-documented capability of CAFs to promote therapy resistance became also apparent following co-cultivation experiments of H2228 and H3122 lung tumor spheroids, as shown by a compromised ability of ALKi treatment to decrease cell proliferation and induce apoptosis. Interestingly, the survival of EML4-ALK v3 driven H2228 cells was consistently better than that of EML4-ALK v1 driven H3128 cells before and after the addition of fibroblasts, which is consistent with the worse outcomes observed for v3-driven ALK+ NSCLC clinical practice (56).

Besides its unprecedented resolution of cellular heterogeneity, scRNA-seq provides a basis for the disentanglement of individual cells in response to stimuli and in the context of their microenvironment (57, 58). In the present study, transcriptionally distinct tumor cell and fibroblast cluster of heterotypic lung tumor spheroids were identified following nonlinear dimensionality reduction using UMAP and cluster marker analysis. The heterogeneity of lung tumor cells could be partially explained by occurring clusters of cycling cells, underlined by congruent expression signatures of identified cluster markers with gene sets utilized for cell cycle annotation (e.g. *TOP2A*, *MKI67*, *PTTG1*, and *CENPF*). The fibroblast population expressed genes related to ECM remodeling, collagen formation, cell adhesion, chemotaxis and other secretory functions, which is consistent with previous reports focusing on the characterization of the tumor microenvironment (59–62). The heterogeneity of CAFs is largely context dependent and increasing evidence points towards functional specializations among CAFs (63). This is highlighted by the identification of different fibroblast populations such as myofibroblastic CAFs (myCAFs) or inflammatory CAFs (iCAFs) (60, 62). Although such stratifications into CAF subtypes could be not delineated herein, a rather mixed phenotype based on partly overlapping gene signatures among different CAF subtypes seemed likely. This finding was suggested to be driven by microenvironmental differences between *in vitro* and *in vivo* condition (e.g. lack of immune compartment or a lymphatic/vascular system) which can substantially influence the fibroblast’s fate (63, 64). Nevertheless, this hypothesis warrants further investigations in the present context.

In oncogene-addicted NSCLC, CAF-derived soluble factors are established to confer treatment resistance through binding to their receptors on the tumor cell’s surface and activating downstream signaling pathways (54, 65). Since the results of CAF-CM promoting the growth of ALK-inhibited tumor spheroids implied a significant contribution of CAF-derived soluble factors towards tumor cell resistance, further analyses focused on the investigation of paracrine signaling pathways as a route of intercellular communication. Among these, HGF represents a CAF-secreted ligand known to induce resistance towards EGFR- and ALK-directed TKIs via stimulation of its cognate receptor MET and downstream PI3K/AKT and MAPK signaling (52, 53). Similarly, a direct link of CAF-derived NRG1 in promoting therapy resistance via ErbB3 receptor signaling was observed in *BRAF* mutant melanoma (38) and in prostate cancer (39). NRG1 was also able to convey therapy insensitivity in NSCLC cells, e.g. through autocrine mechanisms or external stimuli (66–69). In agreement with such observations, the resistance-inducing effect of the CAF-associated ligands HGF and NRG1 on TKI-treated H2228 and H3122 tumor spheroids was confirmed by increased cell viabilities despite ALK inhibition. Contrary to that, the resistance-inducing effects of CAF-CM could be attenuated following the addition of inhibitory antibodies directed against HGF and NRG1.

By analyzing scRNA-seq data, we were able to infer the main molecular drivers of CAF-mediated therapy resistance and uncovered the association of enhanced lipogenic gene expression with a CAF-driven resistant phenotype of ALKi-treated lung adenocarcinoma cells. Activation of *de novo* lipid synthesis is frequently observed in cancer and is assumed to fuel high demands in membrane biogenesis during cell divisions and tumor growth in a cell-autonomous manner (17). Accumulating evidence also suggests that multiple oncogenic signaling pathways converge on lipid synthesis, thereby creating metabolic dependencies of molecularly altered cancers (30, 31, 50, 70). In addition, the induction of SREBP-1-driven *de novo* synthesis of endogenous lipids becomes increasingly apparent in many drug-resistant cancer cells (71) and sustained lipogenesis was shown to be a key mediator of acquired resistance towards moleculary directed drugs in oncogene-driven melanoma (32) and NSCLC (33, 72). Notably, such reprogramming of lipid metabolism is not only influenced by cell-autonomous genomic alterations but also by non-genomic components and non-cell autonomous players, i.e. the tumor microenvironment (73). However, the direct impact of CAFs on inducing lipid pathway activity in therapy-resistant cancer cells, as presented in this project, remains largely unexplored.

Our results further showed that ALK inhibition reduced levels of saturated and mono-unsaturated phospholipids, while increasing lipid poly-unsaturation and lipid peroxidation. Since bis-allylic carbons are particularly reactive, saturated, mono-unsaturated, and polyunsaturated acyl chains differ in their susceptibility to peroxidation by free radicals (51, 74). According to that, the increase in lipid ROS following ALK-TKI treatment aligns with the high unsaturation levels of membrane lipid species. In contrast, presence of CAF-CM largely reversed the lipid composition, rendering ALKi-treated lung tumor spheroids less susceptible to lipid peroxidation by limiting the degree of phospholipid poly-unsaturation. Another study similarly demonstrated that CAFs were able to ameliorate lipid peroxidation product accumulation, accompanied by chemo-resistance induction of gastric cancer cells (75). In recent years, mounting evidence has linked oncogenic RTK activation, downstream signal transduction and metabolic reprogramming in neoplastic cells (76, 77). Previous work demonstrated so far a modulation of SREBP-1 activation via KRAS-driven MAPK activation (50), direct ERK1/2-specific SREBP phosphorylation (78), or mutant *BRAF* (32). Nevertheless, most studies pinpoint to a signal transduction via the PI3K/AKT/mTORC1/SREBP axis downstream of active RTKs to induce *de novo* lipid biosynthesis (25, 28). Our work likewise was able to demonstrate that CAF-CM and the CAF-secreted factors HGF and NRG1 re-induced AKT signaling in ALKi-treated lung tumor spheroids, concomitant with an induction of SREBP-1 and its lipogenic targets. While the in depth roles of CAF-secreted RTK-ligands in regulating tumor cell lipid homeostasis have still not been sufficiently explored, some concordant indications can be deviated from other studies. As such, HGF increased ACSVL3 expression levels in human glioma cells (U373), an enzyme that plays a central role in activating FAs for complex lipid synthesis (79). NRG1 is currently heavily explored for its metabolic role in a variety of tissues (29) and was shown to regulate cholesterol and fatty acid synthesis in the process of Schwann cell myelination (80, 81). Critically, pharmacological targeting of SREBP-1 restored the sensitivity of resistant lung tumor spheroids to lorlatinib and was able to overcome the established CAF-mediated lipid metabolic-supportive niche, thereby pinpointing towards potential new therapeutic options for the clinical treatment of advanced NSCLC. Of note, our findings suggest that these mechanisms are common for both standard (v1) and high-risk (v3) EML4-ALK fusion variants and could therefore provide novel strategies to counter the currently unmet need of ALK+ disease with adverse molecular features (82).

Collectively, the systematic approach of combining advanced 3D cell cultures of *ALK*-translocated NSCLC cell lines and CAFs with scRNA-seq provided insights into disctinct cell (sub)population-specific gene expression variabilities and in response to external stimuli. Our data further established a novel and crucial function of CAFs in facilitating resistance to ALK-TKI treatment through lipid metabolic reprogramming. These findings highlight the significance of CAF-mediated resistances and reveal potential strategies to overcome TKI resistance by targeting vulnerabilities downstream of oncogenic signaling in order to further improve clinical benefit for ALK+ NSCLC patients.

## Materials and Methods

### Chemicals/Reagents

Brigatinib, lorlatinib, and fatostatin were purchased from TargetMol (Boston, MA, USA). TGF-β1 was purchased from PeproTech (Cranbury, NJ). The following compounds were purchased from R&D Systems (Minneapolis, MN, USA): HGF, NRG1-α, NRG1-β1, OPG, and WNT5A. Drug and reagent preparations were conducted according to manufacturer’s instructions. Anti-αSMA (A5228) antibody was obtained from Sigma-Aldrich (St. Louis, MO, USA). Anti-FAP antibody (ab207178) was purchased from Abcam (Cambridge, MA, USA). Anti-Ki-67 antibody, eFluor 450 (48-5698-82) was purchased from (Thermo Fisher Scientific, Waltham, MA, USA). Anti-SREBP-1 antibody (sc-13551) was obtained from SantaCruz Biotechnology (Santa Cruz, CA, USA). Anti-ACLY (#4332), β-Actin (#4970), FASN (#3180), GAPDH (#2118), and secondary horseradish peroxidase (HRP)-conjugated antibodies goat anti-rabbit IgG (#7074) and horse anti-mouse IgG (#7076) were purchased from Cell Signaling Technology (Danvers, MA, USA).

### Cell culture

Human lung cancer cells (A549, NCI-H2228) and fetal lung fibroblasts (MRC-5 [CCL-171]) were purchased from ATCC (Manassas, VA, USA). NCI-H3122 cells were obtained from NCI (Bethesda, MD, USA). Human primary fibroblasts (FB1, FB2) derived from NSCLC adenocarcinoma patients were generated and kindly provided by the Translational Research Unit of the Thoraxklinik (Heidelberg, Germany) following written informed consent and the approval by the ethics committee of the Medical Faculty Heidelberg (S-270/2001, S-296/2016, and S-435/2019, respectively). Shortly, tumor tissues were minced in small pieces and digested using Liberase DH Research Grade (Roche, Mannheim, Germany). Afterwards, cells were filtered using 100 µm and 40 µm cell strainer (Corning, Corning, NY, US) and purified using Histopaque (Sigma Aldrich, St. Louis, MO, US). Cells were seeded in DMEM (Thermo Fisher Scientific) supplemented with 10% FBS (Thermo Fisher Scientific) until fibroblasts reached 80% confluency. NCI-H2228 and NCI-H3122 cells were maintained in RPMI 1640 medium (Thermo Fisher Scientific), MRC-5 cells in EMEM (Thermo Fisher Scientific) and the primary fibroblast lines FB1 and FB2 in DMEM (Thermo Fisher Scientific), all supplemented with 10% FBS (Thermo Fisher Scientific). Cell lines and primary cells were propagated at 37°C in a humidified atmosphere with 5% CO_2_ and no longer maintained than 8-15 passages. Cells were subcultured at 70-80% confluence by harvesting with 0.25% Trypsin-EDTA (Thermo Fisher Scientific), and suspended to required cell density for subsequent assays. Cells were routinely monitored for mycoplasma contamination and authenticated based on single nucleotide polymorphism (SNP)-profiling.

### CAF transdifferentiation

Fibroblasts were seeded 24 h prior to TGF-β1 treatment. On the next day, cells were serum starved for 8 h with growth medium containing 0.5% FCS. Following this, TGF-β1 made up in serum low-medium (final concentration 2 ng/ml) was added to the cell culture vessel and cells were incubated for 72 h. Fibroblasts treated with 0.5% FCS-containing growth medium only served as control.

### 3D cell culture

3D cultures were generated as homo- and heterotypic tumor spheroids. To this end, H2228 and H3122 cells were trypsinized and diluted in ice cold spheroid medium (DMEM supplemented with GlutaMAX + 25 mM HEPES (Thermo Fisher Scientific) + 10% FCS) to. A volume of 50 µl cell suspension corresponding to 2,000 cells was added to each well of a round-bottom ultra-low attachment (ULA) 96-well plate (Corning, NY, USA) followed by centrifugation at 300 × g for 5 min. Adequate spheroid formation was initiated by adding 50 µl of ice-cold spheroid medium additives into corresponding tumor cell-containing wells as follows: H2228 cells were supplemented with 3% growth factor reduced (GFR)-Matrigel (Corning), while H3122 cells were supplemented with 1% GFR-Matrigel and 10 µg/ml Cultrex 3D Culture Matrix Rat Collagen I (Bio-Techne, Minneapolis, MN, USA). The plates were incubated under standard cell culture conditions for 72 h. For the generation of heterotypic spheroids, H2228 and H3122 cells were co-seeded with previously activated MRC-5, FB1 and FB2 fibroblasts, respectively. A volume of 100 µl cell suspension corresponding to 3,000 cells with a tumor cell:fibroblast ratio of 1:6 was added to each well of a round-bottom ULA 96-well plate. Plates were centrifuged at 300 × g for 5 min and incubated under standard cell culture conditions as described above.

Homotypic fibroblast spheroids were generated utilizing the AggreWell 400 24-well plate (Stemcell Technologies, Vancouver, Canada). Plates were prepared according to the manufacturer’s instructions. Previously (non)activated MRC-5, FB1 and FB2 cells were detached and diluted in spheroid medium to 2.88 × 10^7^ cells/ml. A volume of 500 µl cell suspension was seeded into each well containing a microwell inlay corresponding to 1,000 cells/microwell. Fibroblast spheroids were grown under standard cell culture conditions for 48 h.

### CAF-conditioned media

CAF-conditioned media (CAF-CM) were made from TGF-β1-activated FB2 cells grown as fibroblast spheroids as described above. After 48 h, conditioned media were collected and centrifuged for 10 min at 1,000 × g to remove remaining cells and cell debris. The supernatant was immediately stored at -80°C for later usage.

### Spheroid viability

Homotypic spheroids were treated by adding 100 µl of fresh spheroid culture medium supplemented with indicated agents. Cell viability of (non-)treated tumor spheroids was evaluated using the CellTiter-Glo 3D Cell Viability Assay (Promega, Fitchburg, WI, USA) according to the manufacturer’s instructions with slight modifications. Briefly, tumor spheroids were transferred with 50 µl culture medium into a new opaque-walled and white-bottomed 96-well plate and equilibrated to RT. An equivalent volume of reagent was added to all samples and plates vigorously shaken for 5 min at 750 rpm protected from light. Luminescence signals were recorded following incubation for 30 min at RT using the Infinite M200 plate reader (Tecan, Männedorf, Switzerland).

### Spheroid growth

To quantify spheroid size after ALKi treatment w/o and w/ CAF-CM, tumor spheroid mono-cultures were grown for 72 h. On the day of treatment, previously generated CAF-CM was thawed on ice and diluted 1:2 with fresh spheroid medium. H2228 and H3122 spheroids were treated with appropriate ALKi concentrations either in diluted CAF-CM or in spheroid medium only. After 72 h treatment spheroid size was assessed by brightfield imaging at 10x magnification using an Axio Observer 7 (Carl Zeiss, Jena, Germany). Twelve representative spheroids of each corresponding sample group were imaged. Spheroid area (µm²) was evaluated using the freehand selection tool of the image analysis software Fiji (83).

### Magnetic cell separation

Heterotypic spheroids were pooled and dissociated with Accumax - Cell Aggregate Dissociation Medium (Thermo Fisher Scientific) via incubation for 8 min at 37°C, followed by gently pipetting up and down to disrupt cell-cell contacts. Both steps were repeated at least twice until complete dissociation of spheroids and remaining cell clumps were removed by passing the cells through pre-separation filters (20 µm). Tumor cells were separated from the fibroblast population using CD326 (EpCAM) MicroBeads (Miltenyi Biotec, Bergisch Gladbach, Germany) via magnetic-activated cell sorting (MACS) according to the manufacturer’s instructions.

### Annexin V apoptosis assay

Tumor cells were stained with 10 µM CellTrace CFSE Cell Proliferation Kit (Thermo Fisher Scientific) according to the manufacturer’s instructions. Stained cells were resuspended in 1 ml fresh spheroid culture medium and used for homo- and heterotypic spheroid generation as described above. Apoptosis was measured utilizing the Annexin V Apoptosis Detection Kit eFluor 450 assay (Thermo Fisher Scientific) with the Fixable Viability Dye (FVD) eFluor 660 (Thermo Fisher Scientific). Single cells of pooled and dissociated tumor spheroids were suspended in flow cytometry buffer (1X DPBS + 1% BSA) and placed at 37°C for 30 min to allow recovery of the cells. Cells were pelleted by centrifugation for 3 min at 300 × g and recovered in 500 µl FVD-staining solution (FVD diluted 1:1000 in cold 1X DPBS) followed by incubation for 30 min at RT in the dark. Next, cells were processed with Annexin V staining solution according to the manufacturer’s protocol. Samples were run through a FACSCanto II flow cytometry system (BD Biosciences, San Jose, CA, USA) and data analyzed using FlowJo software.

### Cell cycle analysis

After corresponding treatment, pooled and dissociated tumor spheroids were fixed in 70% ice-cold ethanol and stained with anti-Ki-67 eFluor 450 antibody (1:20). Combinatory RNaseA treatment and PI-staining was performed using the FxCycle PI/RNase Staining Solution (Thermo Fisher Scientific). Data of gated tumor cells was acquired by running the samples on a FACSCanto II instrument. Cell cycle plots of quiescent (G0-phase) and actively cycling tumor cells (G1-, S-, G2-M-phase) were generated with FlowJo software.

### Lipid peroxidation quantification by BODIPY C11 staining

Tumor spheroids were pooled, dissociated and stained with 10 µM BODIPY 581/591 C11 (Thermo Fisher Scientific) for 30 minutes at 37°C after indicated treatment. Labeled cells were washed twice with PBS and subjected to flow cytometry analysis using a FACSCanto II instrument (excitation 488 nm). Positive controls were generated by incubating untreated samples with 1 mM H_2_O_2_ for 1 h at 37°C.

### RNA extraction, cDNA synthesis, and qRT-PCR

Total RNA was extracted from cell lines or pooled tumor spheroids using the miRNeasy Mini Kit (Qiagen, Hilden, Germany) following the manufacturer’s protocol. RNA was reverse-transcribed using the RevertAid H Minus First Strand cDNA Synthesis Kit (Thermo Fisher Scientific) according to the supplier’s instructions. Quantitative real-time PCR (qPCR) was carried out in the LightCycler 480 II system (Roche, Basel, Switzerland) using Universal ProbeLibrary (Roche) or primaQUANT (Steinbrenner, Wiesenbach, Germany) SYBR assays with primers listed in Supplementary Table S4. Relative gene expression fold changes were calculated using the 2^(-ΔΔCt)^ method (84), with *ACTB* or *GAPDH* as internal standards.

### Immunoblot analysis

Whole-cell lysates were collected from single cells or pooled tumor spheroids using RIPA buffer, supplemented with protease/phosphatase inhibitor cocktail (Roche). Protein concentration was measured with Pierce™ BCA Protein Assay Kit (Thermo Fisher Scientific) following manufacturer’s instructions. Proteins were separated by SDS-PAGE and transferred onto PVDF membranes applying the Trans-Blot Turbo Transfer system (Bio-Rad Laboratories). Membranes were blocked with 5% BSA or 5% NFDM in 1X TBS-T buffer and incubated with specific primary and secondary antibodies. Development of chemiluminescent immunoblot signals was carried out with the SuperSignal West Dura Extended Duration Substrate (Thermo Fisher Scientific) and detected by a ChemiDoc XRS+ System (Bio-Rad Laboratories).

### CAF secretome profiling

Simultaneous detection and quantification of proteins and growth factor was conducted using a Magnetic Luminex Assay. Analytes in cell culture supernatants of fibroblast spheroids were tested with the Human Premixed Multi-Analyte Kit (Bio-Techne). All reagents, standards, and samples were prepared as recommended by the manufacturer. Data acquisition was conducted using a Bio-Plex 200 system (Bio-Rad Laboratories) and standard curves were created using the five parameter logistic (5-PL) curve-fit. Each standard and sample was measured in duplicates. Secreted levels of WNT5A in cell culture supernatants of fibroblast spheroids were quantitatively determined employing the Human WNT5A ELISA Kit (FineTest, Wuhan, China). All reagents, standards, and samples were prepared according to the manufacturer’s instructions. Absorbance of the samples was recorded at 450 nm using the Infinite M200 plate reader (Tecan). Target concentrations were interpolated from a standard curve created from the standards with predefined concentrations. Each sample was measured in duplicates.

### Single-cell RNA-Sequencing by 10x technology

Single cell suspensions of homo- and heterotypic tumor spheroids were prepared as described before and pipetted through a 20 µm cell strainer. To obtain barcoded scRNA-seq libraries, samples were processed using the Chromium Single Cell 3’ Reagent Kit v3 (10x Genomics, Pleasanton, CA, USA) as per manufacturer’s protocol. In Brief, a total of 10,000 cells were loaded on a Chromium Single-Cell Controller Instrument (10x Genomics) to generate single-cell Gel Bead-in EMulsions (GEMs). Incorporated mRNA was reverse transcribed into barcoded cDNA and PCR-amplified. This was followed by fragmentation, end repair, A-tailing, and indexing to generate sequencing libraries. Enriched scRNA-seq library constructs were subjected to 28+74 bp paired-end sequencing on four lanes on the HiSeq 4000 system (Illumina, San Diego, CA, USA).

### Preprocessing of scRNA data

Single-cell RNA-sequencing data were obtained as raw demultiplexed *.fastq* files and pre-processed using the Cell Ranger (version 3.1.0) analysis pipeline (10x Genomics). In brief, cellular barcodes were demultiplexed and sequencing reads were mapped to the human reference genome (hg38/GRCh38). Aligned reads were filtered and read count matrices obtained by cell barcode and unique molecular identifier (UMI) counting.

### Secondary analysis using Seurat

Further filtering and clustering analyses of pre-processed single-cell RNA data was performed in R (version 3.6.0) using the ‘Seurat v3’ R package (85). Loaded expression matrices were filtered to remove genes detected in <3 cells. Thresholding the number of detected genes and the fraction of mitochondrial reads was performed for quality control measures. Thresholds were determined per dataset by visual inspection of the distributions and low-quality cells were excluded from further analysis. Finally, 66,375 single cells remained for downstream analyses. For all datasets, gene expression values for each cell were normalized to log scale with a scale factor of 10,000. Next, only the top 2,000 genes exhibiting highest variability across cells were considered for further downstream analysis. For integration of multiple datasets (i.e. control vs. treatment, mono-vs. co-culture), anchor correspondences were identified and allowed to run a single integrated analysis on all cells. A linear regression (‘scaling’) was performed prior to dimensional reduction, to eliminate unwanted sources of variation, such as heterogeneity associated with cell cycle stages, and mitochondrial contamination. Subsequently, the scaled data was used as input for PCA, and top significant principal components were estimated using the Elbow plot function. Significant principal components were further used for graph-based clustering into discrete populations. The resolution parameter, which sets indirectly the number of clusters, was empirically evaluated in order to establish discernible clusters with distinctive marker gene expression. Afterwards non-linear dimensional reduction using UMAP was run to project the predicted populations in two dimensions for data visualization and exploitation purposes. To identify marker genes, each cluster was compared to all other cells using the Wilcoxon Rank Sum test for differential expression testing (min.pct = 0.25, logfc.threshold = 0.25). Marker gene expression was visualized using feature plots as well as expression heatmaps. For computing differentially expressed genes between (un)treated homo-versus heterotypic spheroid-derived tumor cells, differential expression testing (Wilcoxon Rank Sum test, min.pct = 0.25, logfc.threshold = 0.25) was performed on all cells while excluding the fibroblast subcluster.

### Gene set enrichment analysis (GSEA)

To investigate perturbed biological processes and pathways upon ALKi and co-cultivation with activated fibroblasts, GSEA (Molecular Signatures Database (MSigDB); http://software.broadinstitute.org/gsea/msigdb; version 4.03.) was conducted on differentially expressed genes between datasets. For GSEA, natural log-expression fold-change values of analyzed genes were used that passed the following thresholds: |log(fold change)| > 0.25 and the gene was detected in > 25% of the cells in each of the compared groups. Enrichment analysis was performed using GSEAPreranked (desktop application v4.0.3) with permutations value set to 1000 and weighted enrichment statistics. Gene sets were pre-filtered according to gene set sizes, with a max of 500 and a min of 10 genes per gene set being allowed. Expression signatures for selected pathways (Hallmarks & Canonical pathways including REACTOME) were downloaded from MSigDB.

### Cell-cell interaction analysis

Physical and intercellular signaling interactions between lung cancer cells and fibroblasts were predicted by applying the R package ‘RNA-magnet’ (version 0.1.0). Prior to analysis, the highly curated list of ligand-receptor pairs (version 1.0.0) was prepared by changing from mouse to human annotations using the biomaRt package (version 2.42.1) and all entries were manually checked. Previously generated Seurat v3 objects were loaded and analyzed including the permitted ligand localizations ‘Secreted’, ‘ECM’ and ‘Membrane’. Accordingly, cellular interactions were scored based on receptors binding to other cell surface molecules, extracellular matrix components or secreted ligands.

### Lipidome profiling by liquid chromatography-mass spectroscopy (LC-MS)

For lipid extraction, pooled and (un)treated tumor spheroids were washed with 154 mM ice-cold ammonium acetate and snap frozen in liquid nitrogen. Spheroids were resuspended in 500 µl MeOH/H2O (80/20, v/v), the suspension was transferred to a glass tube and another 500 µl MeOH/H2O (80/20, v/v) were added. Splash Lipidomix was added as internal standard (10 µl per sample, Avanti Lipids). After addition of 120 µl 0.2 M HCl, 360 µl chloroform, 400 µl chloroform and 400 µl of water with vigorous mixing between the pipetting steps, samples were centrifuged at 3,000 g for 10 min. 700 µl of the lower phase were collected and taken to dryness under a stream of nitrogen gas at 40 °C. Lipids were dissolved in 100 µl isopropanol prior LC-MS analysis. Lipids were separated on a C8 column (Accucore C8 column, 2.6 µm particle size, 50 × 2.1 mm, Thermo Fisher Scientific) mounted on an Ulitmate 3000 HPLC (Thermo Fisher Scientific) and heated to 40°C. The mobile phase buffer A consisted of 0.1% formic acid in CH3CN/H2O (10/90, v/v) and buffer B consisted of 0.1% formic acid in CH3CN/IPOH/H2O (45/45/10, v/v/v). After injection of 3 µl lipid sample, 20% solvent B were maintained for 2 minutes, followed by a linear increase to 99.5% B within 5 minutes, which was maintained for 27 minutes. After returning to 20% B within 1 minutes, the column was re-equilibrated at 20% B for 5 minutes, resulting in a total run time of 40 minutes. The flow rate was maintained at 350 µl/min and the eluent was directed to the ESI source of the QE Plus from 2 to 35 minutes. MS analysis was performed on a Q Exactive Plus mass spectrometer (Thermo Fisher Scientific) applying the following settings:*Scan settings*: Scan range – 200-1600 m/z in full MS mode with switching polarities (neg/pos) and data-dependent fragmentation; Resolution – 70,000, AGC target – 1E6; Max. injection time – 50 ms. *HESI source parameters*: Sheat gas – 30; Aux gas – 10; Sweep gas – 3; Spray voltage – 2.5 kV; Capillary temperature – 320°C; S-lens RF level – 55.0; Aux gas heater temperature – 55°C. *Fragmentation settings:* Resolution – 17,500; AGC target – 1E5; Max. injection time – 50 ms. Peaks corresponding to the calculated lipid masses (±5 ppm) were integrated using El-Maven (https://resources.elucidata.io/elmaven).

### Statistical analysis

Statistical analysis was done using GraphPad Prism 8. Quantitative data was presented as mean ± standard deviation (SD) and all statistical tests were performed of a minimum of three experimental replicates. If not otherwise stated, statistical testing was conducted by two-sided unpaired t-test to determine significant differences between control and corresponding treatment conditions. A p-value of < 0.05 was considered statistically significant (*, p < 0.05; **, p < 0.01; ***, p < 0.001; ****, p < 0.0001).

## Acknowledgements

We would like to thank Jan-Philipp Mallm and the DKFZ Single-Cell Open Lab (scOpenLab) for assistance with the 10x Genomics experiment and the High Throughput Sequencing Group of the DKFZ Genomic Core Facility for providing next-generation sequencing services. We also thank the DKFZ Proteomics Core Facility for providing mass spectrometry services. We are likewise grateful to Sabrina Gerhardt and Simon J. Ogrodnik for excellent technical support and Sabine M. Klauck for critical reading of the article. This research was funded by the German Center for Lung Research (DZL; Grant ID: 82DZL004B4).

## Author contributions

A.D. contributed to conceptualization, methodology, data curation, formal analysis, visualization, writing – original draft, and writing – review and editing. M.A.S. and T.M. contributed to methodology, and writing – review and editing. L.S. contributed to methodology, formal analysis, writing – original draft, and writing – review and editing. U.K. contributed to methodology, and writing – review and editing. A.S., P.C., and M.T. contributed to conceptualization, methodology, and writing – review and editing. H.S. contributed to conceptualization, supervision, methodology, writing – original draft, and writing – review and editing. All authors read and approved the content of the final manuscript.

## References

1. Soda M, Choi YL, Enomoto M, Takada S, Yamashita Y, Ishikawa S, et al. Identification of the transforming EML4-ALK fusion gene in non-small-cell lung cancer. Nature. 2007;448(7153):561–6.

2. Hallberg B, Palmer RH. The role of the ALK receptor in cancer biology. Ann Oncol. 2016;27 Suppl 3:iii4–iii15.

3. Yuan M, Huang LL, Chen JH, Wu J, Xu Q. The emerging treatment landscape of targeted therapy in non-small-cell lung cancer. Signal Transduct Target Ther. 2019;4:61.

4. Elsayed M, Christopoulos P. Therapeutic Sequencing in ALK(+) NSCLC. Pharmaceuticals (Basel). 2021;14(2).

5. Katayama R, Lovly CM, Shaw AT. Therapeutic targeting of anaplastic lymphoma kinase in lung cancer: a paradigm for precision cancer medicine. Clin Cancer Res. 2015;21(10):2227–35.

6. Lin JJ, Riely GJ, Shaw AT. Targeting ALK: Precision Medicine Takes on Drug Resistance. Cancer Discov. 2017;7(2):137–55.

7. Yoda S, Lin JJ, Lawrence MS, Burke BJ, Friboulet L, Langenbucher A, et al. Sequential ALK Inhibitors Can Select for Lorlatinib-Resistant Compound ALK Mutations in ALK-Positive Lung Cancer. Cancer Discov. 2018;8(6):714–29.

8. Valkenburg KC, de Groot AE, Pienta KJ. Targeting the tumour stroma to improve cancer therapy. Nat Rev Clin Oncol. 2018;15(6):366–81.

9. Altorki NK, Markowitz GJ, Gao D, Port JL, Saxena A, Stiles B, et al. The lung microenvironment: an important regulator of tumour growth and metastasis. Nat Rev Cancer. 2019;19(1):9–31.

10. Zeltz C, Primac I, Erusappan P, Alam J, Noel A, Gullberg D. Cancer-associated fibroblasts in desmoplastic tumors: emerging role of integrins. Semin Cancer Biol. 2020;62:166–81.

11. Wu F, Yang J, Liu J, Wang Y, Mu J, Zeng Q, et al. Signaling pathways in cancer-associated fibroblasts and targeted therapy for cancer. Signal Transduct Target Ther. 2021;6(1):218.

12. Monteran L, Erez N. The Dark Side of Fibroblasts: Cancer-Associated Fibroblasts as Mediators of Immunosuppression in the Tumor Microenvironment. Front Immunol. 2019;10:1835.

13. Lyssiotis CA, Kimmelman AC. Metabolic Interactions in the Tumor Microenvironment. Trends Cell Biol. 2017;27(11):863–75.

14. Hu H, Piotrowska Z, Hare PJ, Chen H, Mulvey HE, Mayfield A, et al. Three subtypes of lung cancer fibroblasts define distinct therapeutic paradigms. Cancer Cell. 2021;39(11):1531–47 e10.

15. Su S, Chen J, Yao H, Liu J, Yu S, Lao L, et al. CD10(+)GPR77(+) Cancer-Associated Fibroblasts Promote Cancer Formation and Chemoresistance by Sustaining Cancer Stemness. Cell. 2018;172(4):841–56 e16.

16. Hanahan D, Weinberg RA. Hallmarks of cancer: the next generation. Cell. 2011;144(5):646–74.

17. Rohrig F, Schulze A. The multifaceted roles of fatty acid synthesis in cancer. Nat Rev Cancer. 2016;16(11):732–49.

18. Beloribi-Djefaflia S, Vasseur S, Guillaumond F. Lipid metabolic reprogramming in cancer cells. Oncogenesis. 2016;5(1):e189.

19. Menendez JA, Lupu R. Fatty acid synthase and the lipogenic phenotype in cancer pathogenesis. Nat Rev Cancer. 2007;7(10):763–77.

20. Griffiths B, Lewis CA, Bensaad K, Ros S, Zhang Q, Ferber EC, et al. Sterol regulatory element binding protein-dependent regulation of lipid synthesis supports cell survival and tumor growth. Cancer Metab. 2013;1(1):3.

21. Zhao Q, Lin X, Wang G. Targeting SREBP-1-Mediated Lipogenesis as Potential Strategies for Cancer. Front Oncol. 2022;12:952371.

22. Shimano H, Sato R. SREBP-regulated lipid metabolism: convergent physiology - divergent pathophysiology. Nat Rev Endocrinol. 2017;13(12):710–30.

23. Shimano H. SREBPs: physiology and pathophysiology of the SREBP family. FEBS J. 2009;276(3):616–21.

24. Jiang T, Zhang G, Lou Z. Role of the Sterol Regulatory Element Binding Protein Pathway in Tumorigenesis. Front Oncol. 2020;10:1788.

25. Hoxhaj G, Manning BD. The PI3K-AKT network at the interface of oncogenic signalling and cancer metabolism. Nat Rev Cancer. 2020;20(2):74–88.

26. Yecies JL, Zhang HH, Menon S, Liu S, Yecies D, Lipovsky AI, et al. Akt stimulates hepatic SREBP1c and lipogenesis through parallel mTORC1-dependent and independent pathways. Cell Metab. 2011;14(1):21–32.

27. Koundouros N, Poulogiannis G. Reprogramming of fatty acid metabolism in cancer. Br J Cancer. 2020;122(1):4–22.

28. Porstmann T, Santos CR, Griffiths B, Cully M, Wu M, Leevers S, et al. SREBP activity is regulated by mTORC1 and contributes to Akt-dependent cell growth. Cell Metab. 2008;8(3):224–36.

29. Zhao M, Jung Y, Jiang Z, Svensson KJ. Regulation of Energy Metabolism by Receptor Tyrosine Kinase Ligands. Front Physiol. 2020;11:354.

30. Bi J, Ichu TA, Zanca C, Yang H, Zhang W, Gu Y, et al. Oncogene Amplification in Growth Factor Signaling Pathways Renders Cancers Dependent on Membrane Lipid Remodeling. Cell Metab. 2019;30(3):525–38 e8.

31. Gouw AM, Margulis K, Liu NS, Raman SJ, Mancuso A, Toal GG, et al. The MYC Oncogene Cooperates with Sterol-Regulated Element-Binding Protein to Regulate Lipogenesis Essential for Neoplastic Growth. Cell Metab. 2019;30(3):556–72 e5.

32. Talebi A, Dehairs J, Rambow F, Rogiers A, Nittner D, Derua R, et al. Sustained SREBP-1-dependent lipogenesis as a key mediator of resistance to BRAF-targeted therapy. Nat Commun. 2018;9(1):2500.

33. Chen Z, Yu D, Owonikoko TK, Ramalingam SS, Sun SY. Induction of SREBP1 degradation coupled with suppression of SREBP1-mediated lipogenesis impacts the response of EGFR mutant NSCLC cells to osimertinib. Oncogene. 2021;40(49):6653–65.

34. Baccin C, Al-Sabah J, Velten L, Helbling PM, Grunschlager F, Hernandez-Malmierca P, et al. Combined single-cell and spatial transcriptomics reveal the molecular, cellular and spatial bone marrow niche organization. Nat Cell Biol. 2020;22(1):38–48.

35. Ding X, Ji J, Jiang J, Cai Q, Wang C, Shi M, et al. HGF-mediated crosstalk between cancer-associated fibroblasts and MET-unamplified gastric cancer cells activates coordinated tumorigenesis and metastasis. Cell Death Dis. 2018;9(9):867.

36. Ko B, He T, Gadgeel S, Halmos B. MET/HGF pathway activation as a paradigm of resistance to targeted therapies. Ann Transl Med. 2017;5(1):4.

37. Lopez-Bergami P, Barbero G. The emerging role of Wnt5a in the promotion of a pro-inflammatory and immunosuppressive tumor microenvironment. Cancer Metastasis Rev. 2020;39(3):933–52.

38. Capparelli C, Rosenbaum S, Berger AC, Aplin AE. Fibroblast-derived neuregulin 1 promotes compensatory ErbB3 receptor signaling in mutant BRAF melanoma. J Biol Chem. 2015;290(40):24267–77.

39. Zhang Z, Karthaus WR, Lee YS, Gao VR, Wu C, Russo JW, et al. Tumor Microenvironment-Derived NRG1 Promotes Antiandrogen Resistance in Prostate Cancer. Cancer Cell. 2020;38(2):279–96 e9.

40. Michailov GV, Sereda MW, Brinkmann BG, Fischer TM, Haug B, Birchmeier C, et al. Axonal neuregulin-1 regulates myelin sheath thickness. Science. 2004;304(5671):700-3.

41. Taveggia C, Zanazzi G, Petrylak A, Yano H, Rosenbluth J, Einheber S, et al. Neuregulin-1 type III determines the ensheathment fate of axons. Neuron. 2005;47(5):681–94.

42. Taveggia C, Thaker P, Petrylak A, Caporaso GL, Toews A, Falls DL, et al. Type III neuregulin-1 promotes oligodendrocyte myelination. Glia. 2008;56(3):284–93.

43. Lane D, Matte I, Rancourt C, Piche A. Osteoprotegerin (OPG) protects ovarian cancer cells from TRAIL-induced apoptosis but does not contribute to malignant ascites-mediated attenuation of TRAIL-induced apoptosis. J Ovarian Res. 2012;5(1):34.

44. De Toni EN, Thieme SE, Herbst A, Behrens A, Stieber P, Jung A, et al. OPG is regulated by beta-catenin and mediates resistance to TRAIL-induced apoptosis in colon cancer. Clin Cancer Res. 2008;14(15):4713–8.

45. Falls DL. Neuregulins: functions, forms, and signaling strategies. Exp Cell Res. 2003;284(1):14–30.

46. Harrison P, Bradley L, Bomford A. Mechanism of regulation of HGF/SF gene expression in fibroblasts by TGF-beta1. Biochem Biophys Res Commun. 2000;271(1):203–11.

47. Matsumoto K, Tajima H, Okazaki H, Nakamura T. Negative regulation of hepatocyte growth factor gene expression in human lung fibroblasts and leukemic cells by transforming growth factor-beta 1 and glucocorticoids. J Biol Chem. 1992;267(35):24917–20.

48. Chiarle R, Voena C, Ambrogio C, Piva R, Inghirami G. The anaplastic lymphoma kinase in the pathogenesis of cancer. Nat Rev Cancer. 2008;8(1):11–23.

49. Yi J, Zhu J, Wu J, Thompson CB, Jiang X. Oncogenic activation of PI3K-AKT-mTOR signaling suppresses ferroptosis via SREBP-mediated lipogenesis. Proc Natl Acad Sci U S A. 2020;117(49):31189–97.

50. Ricoult SJ, Yecies JL, Ben-Sahra I, Manning BD. Oncogenic PI3K and K-Ras stimulate de novo lipid synthesis through mTORC1 and SREBP. Oncogene. 2016;35(10):1250–60.

51. Rysman E, Brusselmans K, Scheys K, Timmermans L, Derua R, Munck S, et al. De novo lipogenesis protects cancer cells from free radicals and chemotherapeutics by promoting membrane lipid saturation. Cancer Res. 2010;70(20):8117–26.

52. Wang W, Li Q, Yamada T, Matsumoto K, Matsumoto I, Oda M, et al. Crosstalk to stromal fibroblasts induces resistance of lung cancer to epidermal growth factor receptor tyrosine kinase inhibitors. Clin Cancer Res. 2009;15(21):6630–8.

53. Yamada T, Takeuchi S, Nakade J, Kita K, Nakagawa T, Nanjo S, et al. Paracrine receptor activation by microenvironment triggers bypass survival signals and ALK inhibitor resistance in EML4-ALK lung cancer cells. Clin Cancer Res. 2012;18(13):3592–602.

54. Shien K, Papadimitrakopoulou VA, Ruder D, Behrens C, Shen L, Kalhor N, et al. JAK1/STAT3 Activation through a Proinflammatory Cytokine Pathway Leads to Resistance to Molecularly Targeted Therapy in Non-Small Cell Lung Cancer. Mol Cancer Ther. 2017;16(10):2234–45.

55. Remsing Rix LL, Sumi NJ, Hu Q, Desai B, Bryant AT, Li X, et al. IGF-binding proteins secreted by cancer-associated fibroblasts induce context-dependent drug sensitization of lung cancer cells. Sci Signal. 2022;15(747):eabj5879.

56. Christopoulos P, Kirchner M, Endris V, Stenzinger A, Thomas M. EML4-ALK V3, treatment resistance, and survival: refining the diagnosis of ALK(+) NSCLC. J Thorac Dis. 2018;10(Suppl 17):S1989–S91.

57. Hwang B, Lee JH, Bang D. Single-cell RNA sequencing technologies and bioinformatics pipelines. Exp Mol Med. 2018;50(8):96.

58. Stuart T, Satija R. Integrative single-cell analysis. Nat Rev Genet. 2019;20(5):257–72.

59. Bartoschek M, Oskolkov N, Bocci M, Lovrot J, Larsson C, Sommarin M, et al. Spatially and functionally distinct subclasses of breast cancer-associated fibroblasts revealed by single cell RNA sequencing. Nat Commun. 2018;9(1):5150.

60. Chen Z, Zhou L, Liu L, Hou Y, Xiong M, Yang Y, et al. Single-cell RNA sequencing highlights the role of inflammatory cancer-associated fibroblasts in bladder urothelial carcinoma. Nat Commun. 2020;11(1):5077.

61. Lambrechts D, Wauters E, Boeckx B, Aibar S, Nittner D, Burton O, et al. Phenotype molding of stromal cells in the lung tumor microenvironment. Nat Med. 2018;24(8):1277–89.

62. Ohlund D, Handly-Santana A, Biffi G, Elyada E, Almeida AS, Ponz-Sarvise M, et al. Distinct populations of inflammatory fibroblasts and myofibroblasts in pancreatic cancer. J Exp Med. 2017;214(3):579–96.

63. Kalluri R. The biology and function of fibroblasts in cancer. Nat Rev Cancer. 2016;16(9):582–98.

64. Sahai E, Astsaturov I, Cukierman E, DeNardo DG, Egeblad M, Evans RM, et al. A framework for advancing our understanding of cancer-associated fibroblasts. Nat Rev Cancer. 2020;20(3):174–86.

65. Yi Y, Zeng S, Wang Z, Wu M, Ma Y, Ye X, et al. Cancer-associated fibroblasts promote epithelial-mesenchymal transition and EGFR-TKI resistance of non-small cell lung cancers via HGF/IGF-1/ANXA2 signaling. Biochim Biophys Acta Mol Basis Dis. 2018;1864(3):793–803.

66. Dong X, Fernandez-Salas E, Li E, Wang S. Elucidation of Resistance Mechanisms to Second-Generation ALK Inhibitors Alectinib and Ceritinib in Non-Small Cell Lung Cancer Cells. Neoplasia. 2016;18(3):162–71.

67. Hegde GV, de la Cruz CC, Chiu C, Alag N, Schaefer G, Crocker L, et al. Blocking NRG1 and other ligand-mediated Her4 signaling enhances the magnitude and duration of the chemotherapeutic response of non-small cell lung cancer. Sci Transl Med. 2013;5(171):171ra18.

68. Isozaki H, Ichihara E, Takigawa N, Ohashi K, Ochi N, Yasugi M, et al. Non-Small Cell Lung Cancer Cells Acquire Resistance to the ALK Inhibitor Alectinib by Activating Alternative Receptor Tyrosine Kinases. Cancer Res. 2016;76(6):1506–16.

69. Kimura M, Endo H, Inoue T, Nishino K, Uchida J, Kumagai T, et al. Analysis of ERBB ligand-induced resistance mechanism to crizotinib by primary culture of lung adenocarcinoma with EML4-ALK fusion gene. J Thorac Oncol. 2015;10(3):527–30.

70. Shackelford DB, Shaw RJ. The LKB1-AMPK pathway: metabolism and growth control in tumour suppression. Nat Rev Cancer. 2009;9(8):563–75.

71. Bacci M, Lorito N, Smiriglia A, Morandi A. Fat and Furious: Lipid Metabolism in Antitumoral Therapy Response and Resistance. Trends Cancer. 2021;7(3):198–213.

72. Xu C, Zhang L, Wang D, Jiang S, Cao D, Zhao Z, et al. Lipidomics reveals that sustained SREBP-1-dependent lipogenesis is a key mediator of gefitinib-acquired resistance in EGFR-mutant lung cancer. Cell Death Discov. 2021;7(1):353.

73. Cai L, Ying M, Wu H. Microenvironmental Factors Modulating Tumor Lipid Metabolism: Paving the Way to Better Antitumoral Therapy. Front Oncol. 2021;11:777273.

74. Reis A, Spickett CM. Chemistry of phospholipid oxidation. Biochim Biophys Acta. 2012;1818(10):2374–87.

75. Zhang H, Deng T, Liu R, Ning T, Yang H, Liu D, et al. CAF secreted miR-522 suppresses ferroptosis and promotes acquired chemo-resistance in gastric cancer. Mol Cancer. 2020;19(1):43.

76. Jin N, Bi A, Lan X, Xu J, Wang X, Liu Y, et al. Identification of metabolic vulnerabilities of receptor tyrosine kinases-driven cancer. Nat Commun. 2019;10(1):2701.

77. DeBerardinis RJ, Chandel NS. Fundamentals of cancer metabolism. Sci Adv. 2016;2(5):e1600200.

78. Roth G, Kotzka J, Kremer L, Lehr S, Lohaus C, Meyer HE, et al. MAP kinases Erk1/2 phosphorylate sterol regulatory element-binding protein (SREBP)-1a at serine 117 in vitro. J Biol Chem. 2000;275(43):33302–7.

79. Pei Z, Sun P, Huang P, Lal B, Laterra J, Watkins PA. Acyl-CoA synthetase VL3 knockdown inhibits human glioma cell proliferation and tumorigenicity. Cancer Res. 2009;69(24):9175–82.

80. D’Antonio M, Musner N, Scapin C, Ungaro D, Del Carro U, Ron D, et al. Resetting translational homeostasis restores myelination in Charcot-Marie-Tooth disease type 1B mice. J Exp Med. 2013;210(4):821–38.

81. Haskins JW, Zhang S, Means RE, Kelleher JK, Cline GW, Canfran-Duque A, et al. Neuregulin-activated ERBB4 induces the SREBP-2 cholesterol biosynthetic pathway and increases low-density lipoprotein uptake. Sci Signal. 2015;8(401):ra111.

82. Christopoulos P, Kirchner M, Bozorgmehr F, Endris V, Elsayed M, Budczies J, et al. Identification of a highly lethal V3(+) TP53(+) subset in ALK(+) lung adenocarcinoma. Int J Cancer. 2019;144(1):190–9.

83. Schindelin J, Arganda-Carreras I, Frise E, Kaynig V, Longair M, Pietzsch T, et al. Fiji: an open-source platform for biological-image analysis. Nat Methods. 2012;9(7):676-82.

84. Pfaffl MW. A new mathematical model for relative quantification in real-time RT-PCR. Nucleic Acids Res. 2001;29(9):e45.

85. Butler A, Hoffman P, Smibert P, Papalexi E, Satija R. Integrating single-cell transcriptomic data across different conditions, technologies, and species. Nat Biotechnol. 2018;36(5):411–20.

