## Supplementary Information for "Cancer-associated fibroblasts promote drug resistance in *ALK*-driven lung adenocarcinoma cells by upregulating lipid biosynthesis"

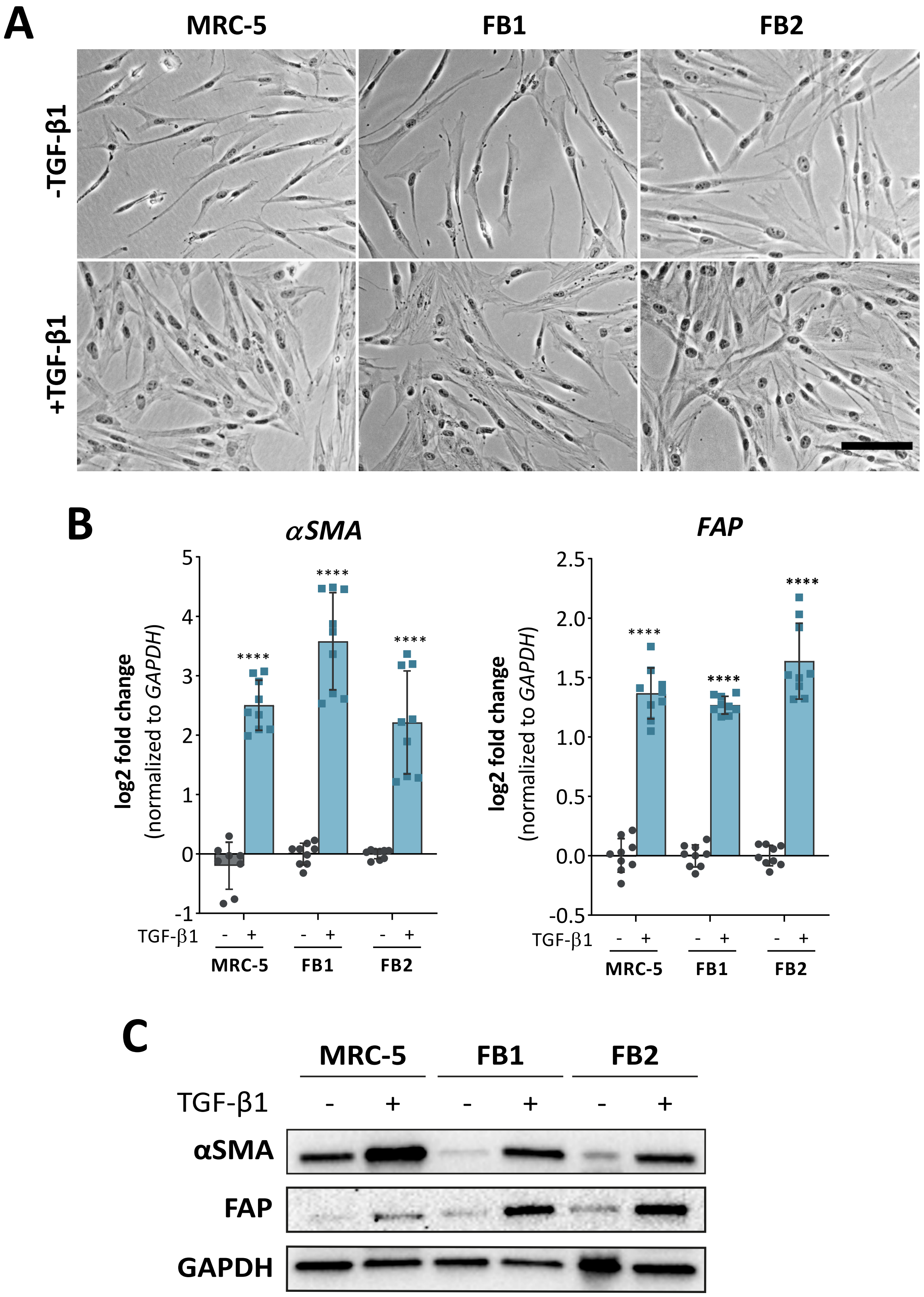


***Supplementary Figure S1:*** (A) Phase-contrast images of untreated (top) versus transforming growth factor-beta 1 (TGF-β1)-stimulated (bottom) MRC-5 fibroblasts and two primary fibroblast lines (FB1, FB2). Scale bar: 100 µm. Validation of CAF transformation 72h following TGF-β1 treatment as shown by expression changes of alpha-smooth muscle actin (αSMA) and fibroblast-activation protein (FAP) on mRNA (B) and protein level (C) (n = 3). Data are presented as mean ± SD. ****, $p \leq 0.0001$.

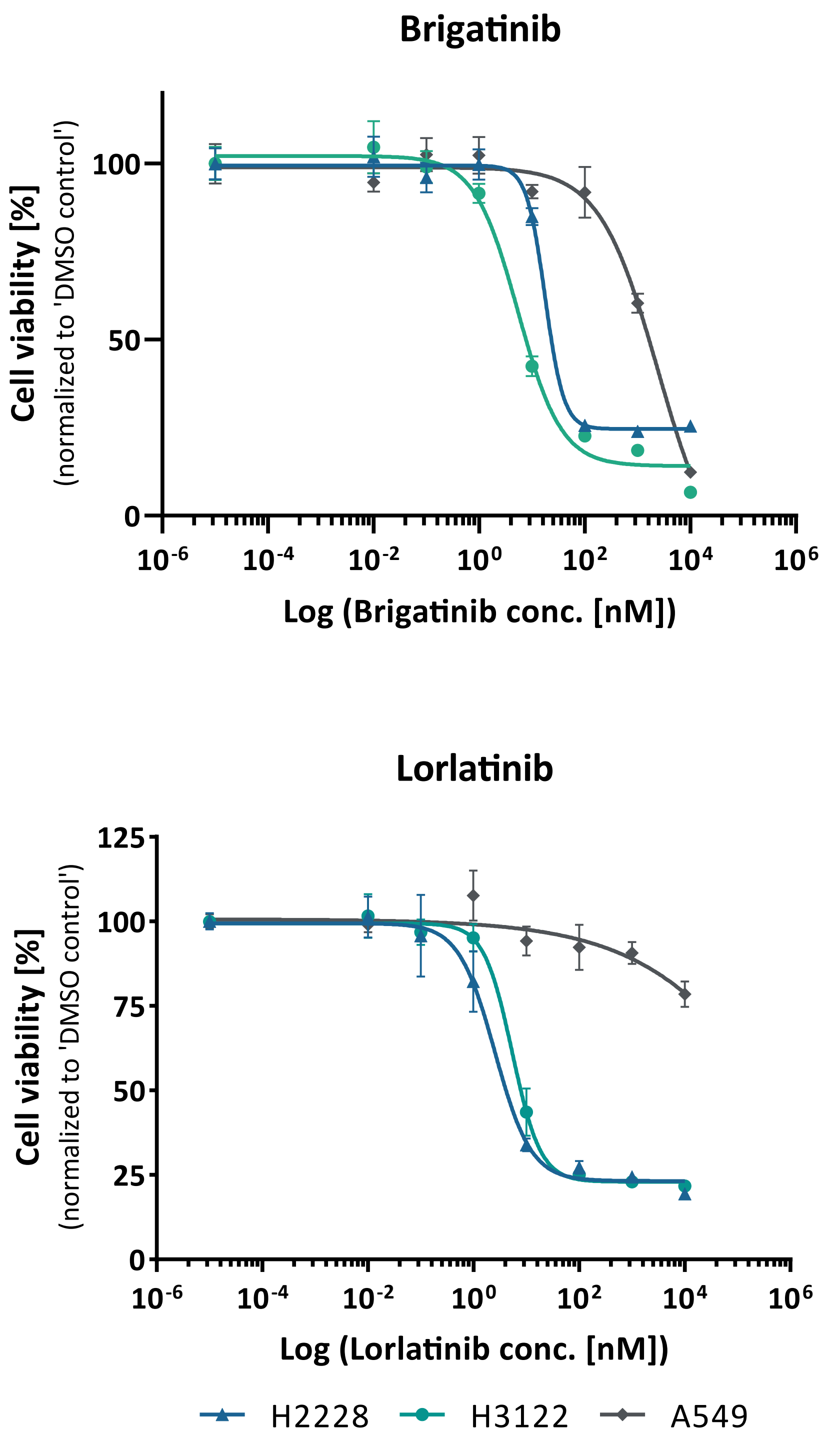


***Supplementary Figure S2:*** Representative dose-response curves of human NSCLC cell lines H2228, H3122, and A549 following 72 h of treatment with the ALK-TKIs brigatinib and lorlatinib.

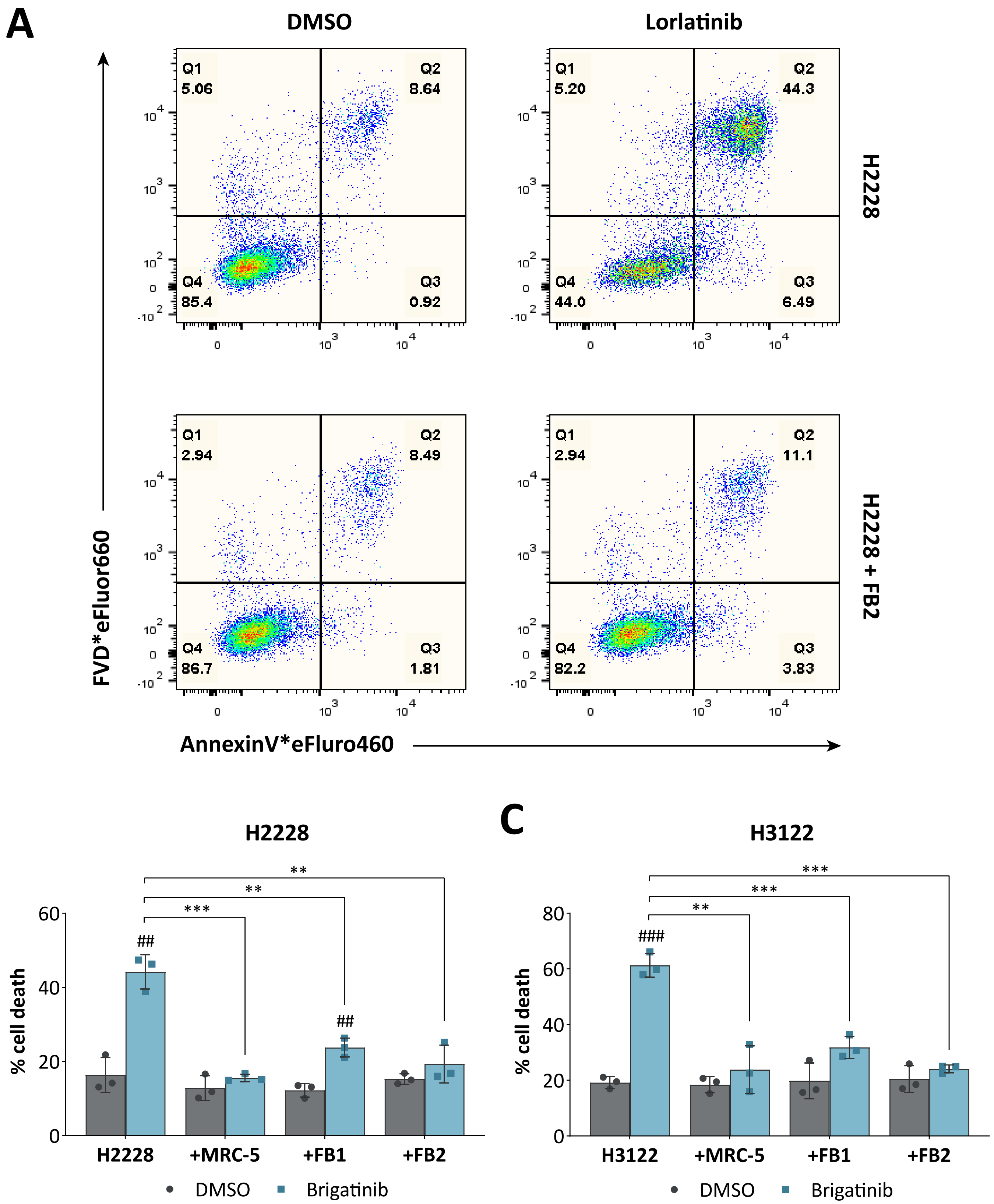


***Supplementary Figure S3:*** (A) Representative dot plots showing the amount of dead or dying H2228 cells derived from dissociated (non-)treated homo- and heterotypic tumor spheroids following flow cytometric cell death analysis. Quantification of cell death rates of H2228 (B) and H3122 (C) cells as given by the sum of early and late apoptotic/necrotic cells (n = 3). Data are presented as mean ± SD. ## ,p≤0.01; ###, p≤0.001 compared to corresponding DMSO controls. **, p≤0.01; ***, p≤0.001 in comparison to brigatinib-treated mono-cultures. FVD, fixable viability dye.

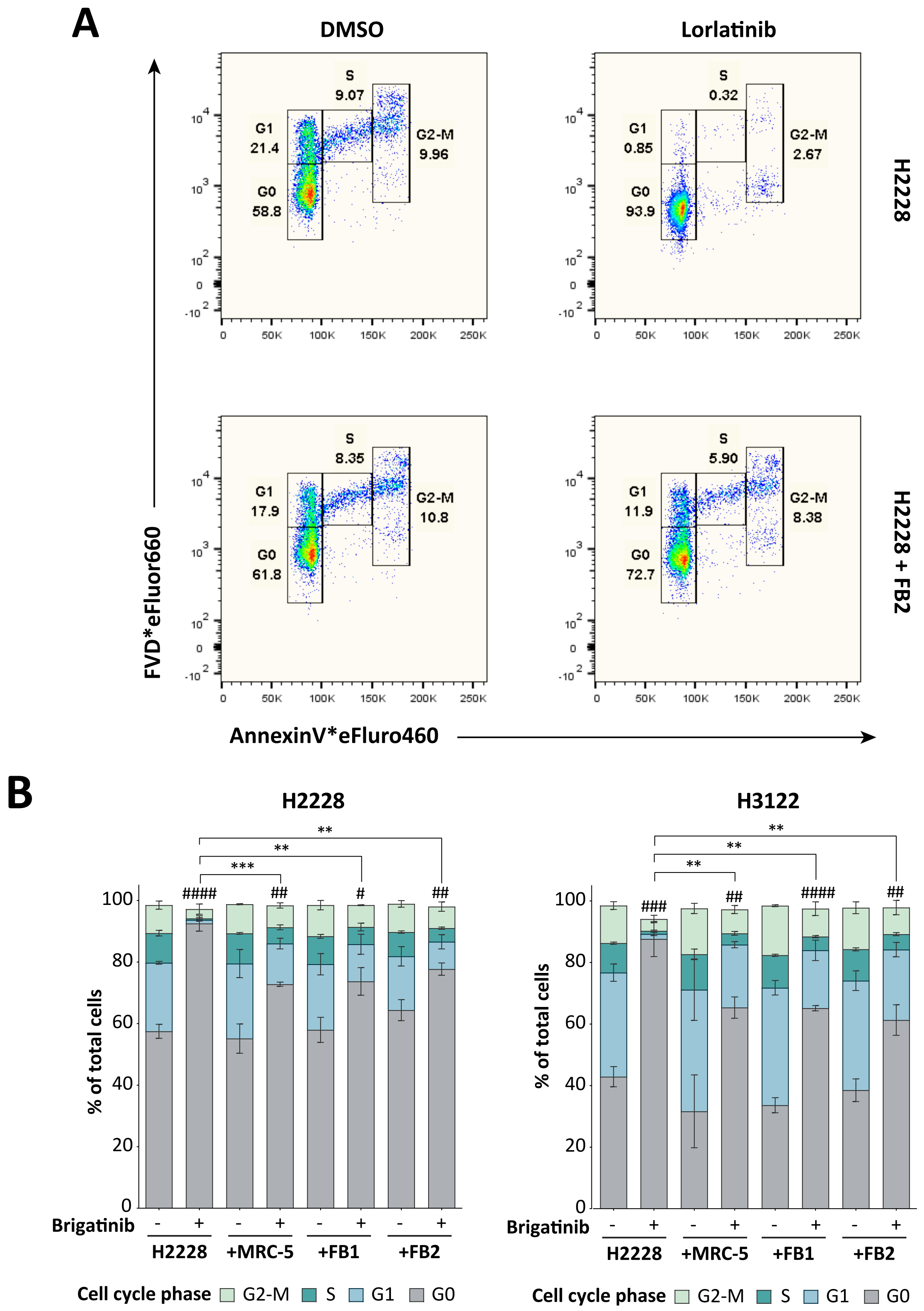


***Supplementary Figure S4:*** (A) Representative dot plots illustrating the distribution among the G0, G1, S, and G2-M cell cycle phases of H2228 cells derived from dissociated (non-)treated homo- and heterotypic tumor spheroids following flow cytometric cell cycle analysis. Quantification of the portion of H2228 (B) and H3122 (C) cells according to their cell cycle phase status (n = 3). Data are presented as mean ± SD. #,p≤0.05; ##, p≤0.01; ###, p≤0.001; ####, p≤0.0001 of cells wihtin the G0-phase compared to corresponding DMSO controls. **, p≤0.01; ***, p≤0.001 of cells wihtin the G0-phase in comparison to brigatinib-treated mono-cultures. PI, propidium iodide.

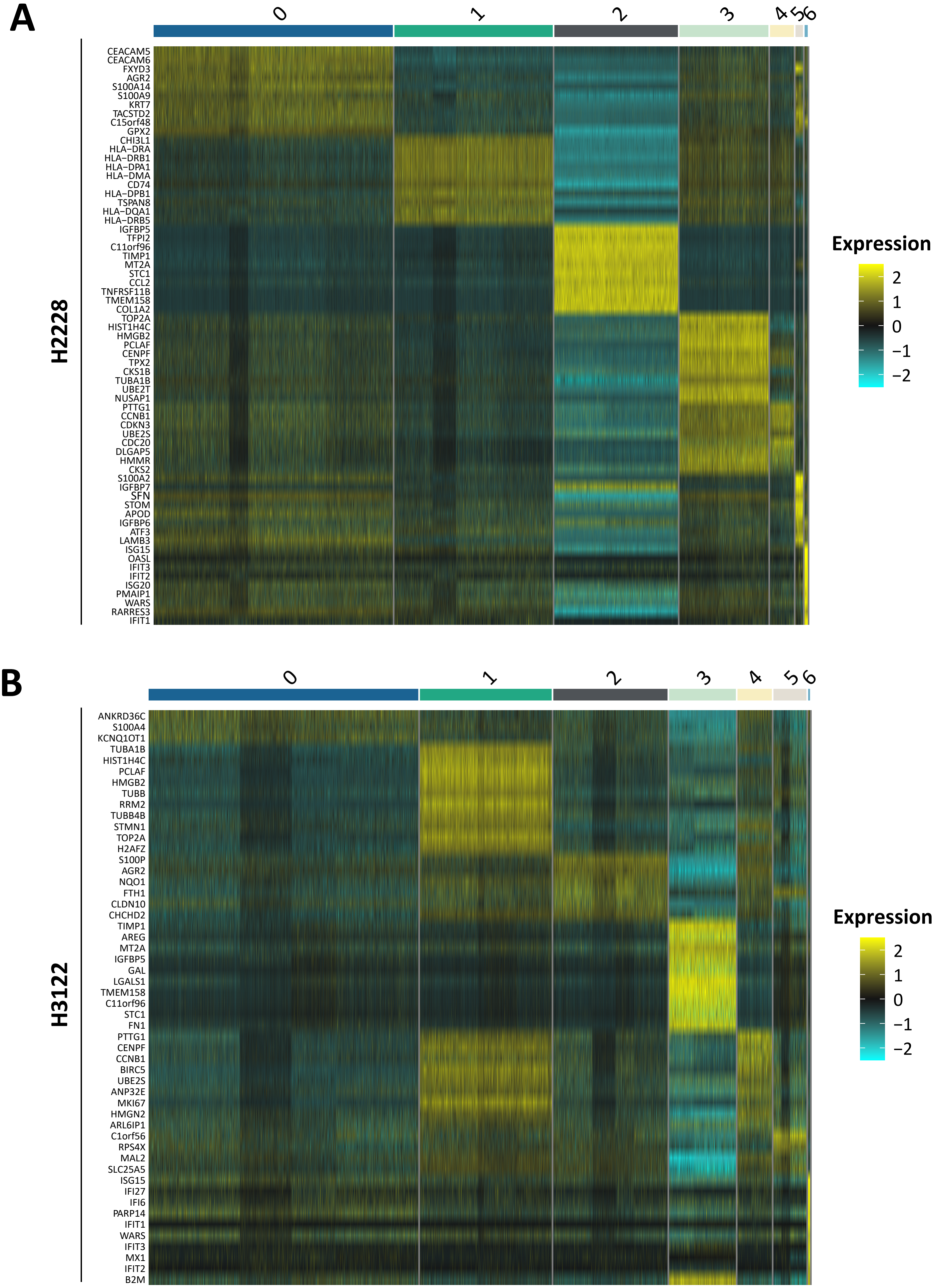


***Supplementary Figure S5:*** Heatmaps depict top ten significantly expressed marker genes of each cell cluster identified following clustering analysis of H2228 (A) and H3122 (B) samples.

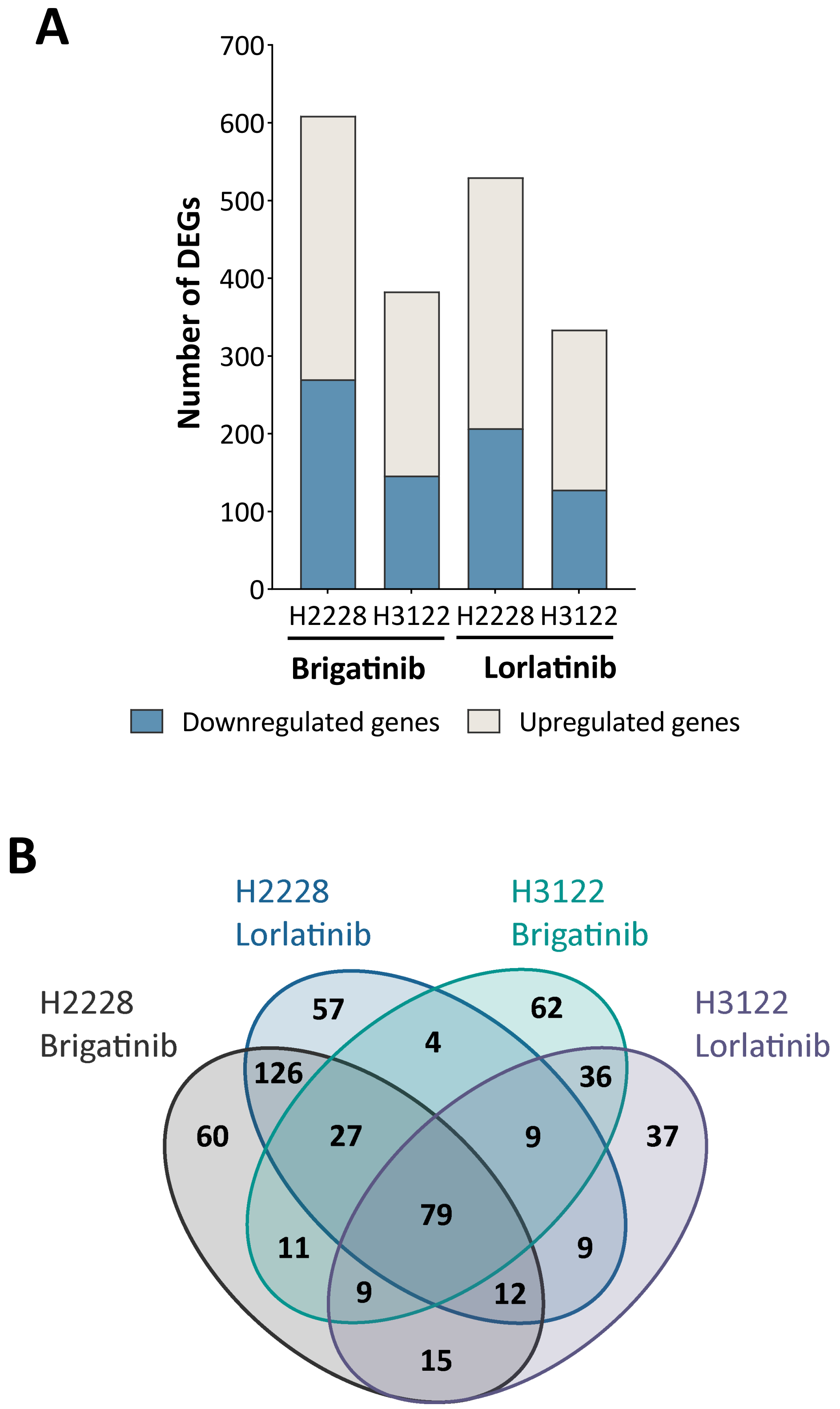


***Supplementary Figure S6:*** (A) Bar graphs depict the total number of differentially expressed genes (DEGs) observed between ALK-inhibited mono-cultured and FB2 co-cultured H2228 and H3122 cells. Blue bars represent number of downregulated genes, while green bars represent upregulated genes. (B) Venn diagram illustrates the DEGs upregulated in FB2 co-cultures versus mono-cultures between brigatinib- or lorlatinib-treated H2228 and H3122 cells.

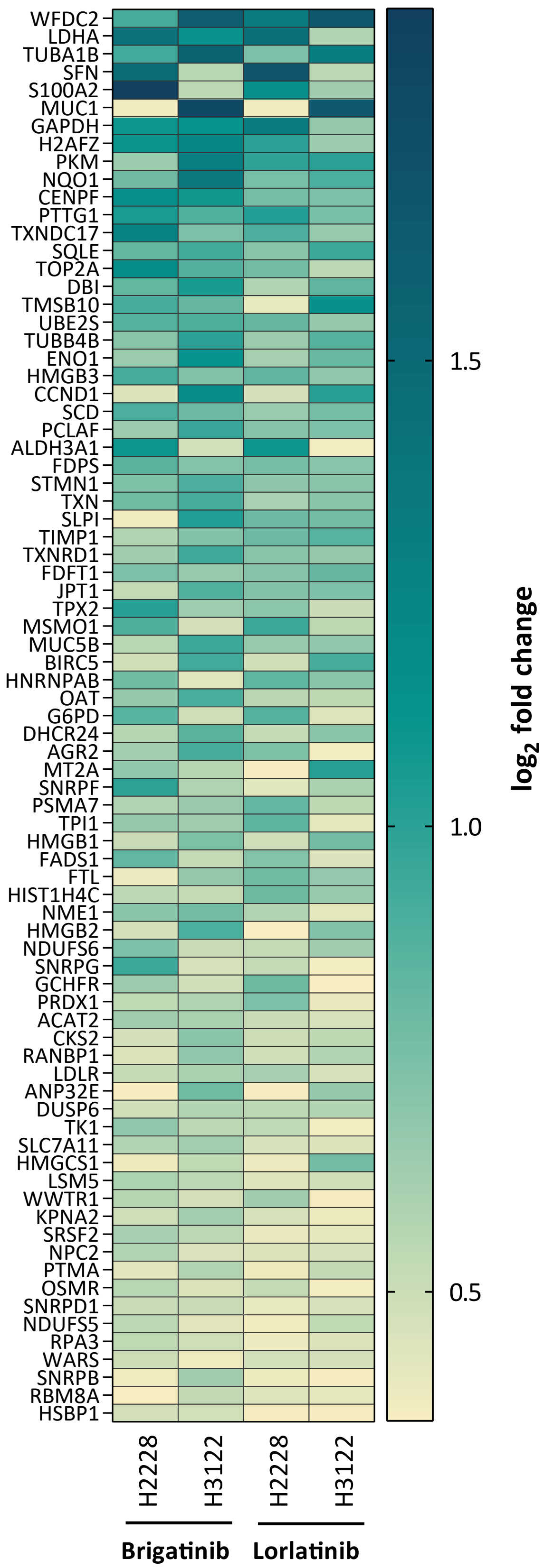


***Supplementary Figure S7:*** Heatmap of differentially expressed genes upon ALK inhibition. The set of 79 upregulated genes overlapping between brigatinib- and lorlatinib-treated H2228 and H3122 co-cultures in comparison to mono-cultures are color-coded according to their log2 fold-change expression values.

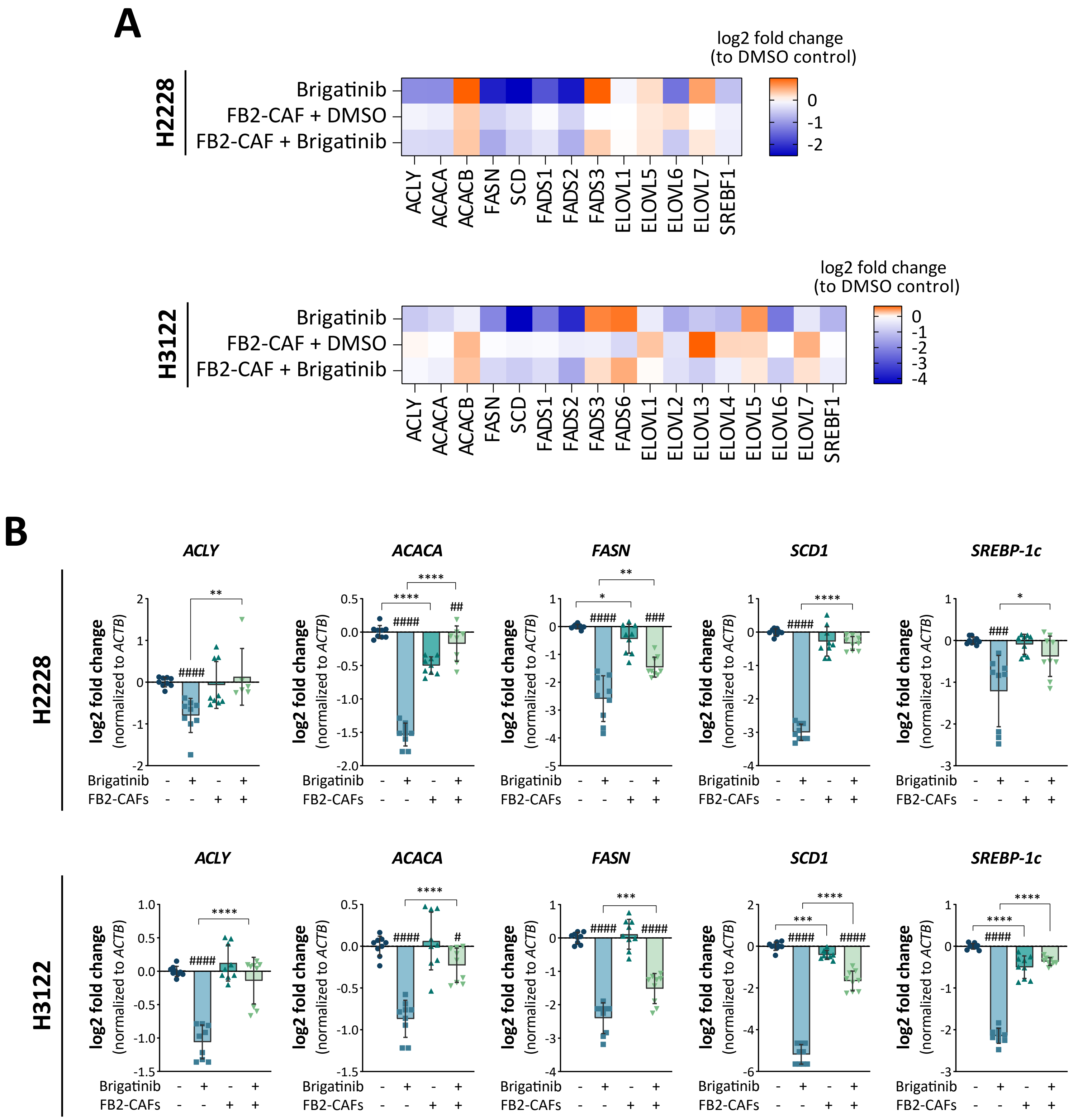


***Supplementary Figure S8:*** (A) Expression heatmap of fatty acid metabolism-related genes in single-cell transcriptome datasets of brigatinib-treated H2228 and H3122 cells. (B) Co-cultivation with FB2-CAFs influences expression of fatty acid metabolism-related targets upon ALK signaling perturbation using brigatinib in H2228 and H3122 cells (n = 3). Data are presented as mean ± SD. #, p≤0.05; ##, p≤0.01; ####, p≤0.0001 compared to corresponding DMSO controls. *, p≤0.05; **, p≤0.01; ***, p≤0.001; ****, p≤0.0001 in comparison to brigatinib-treated mono-cultures.

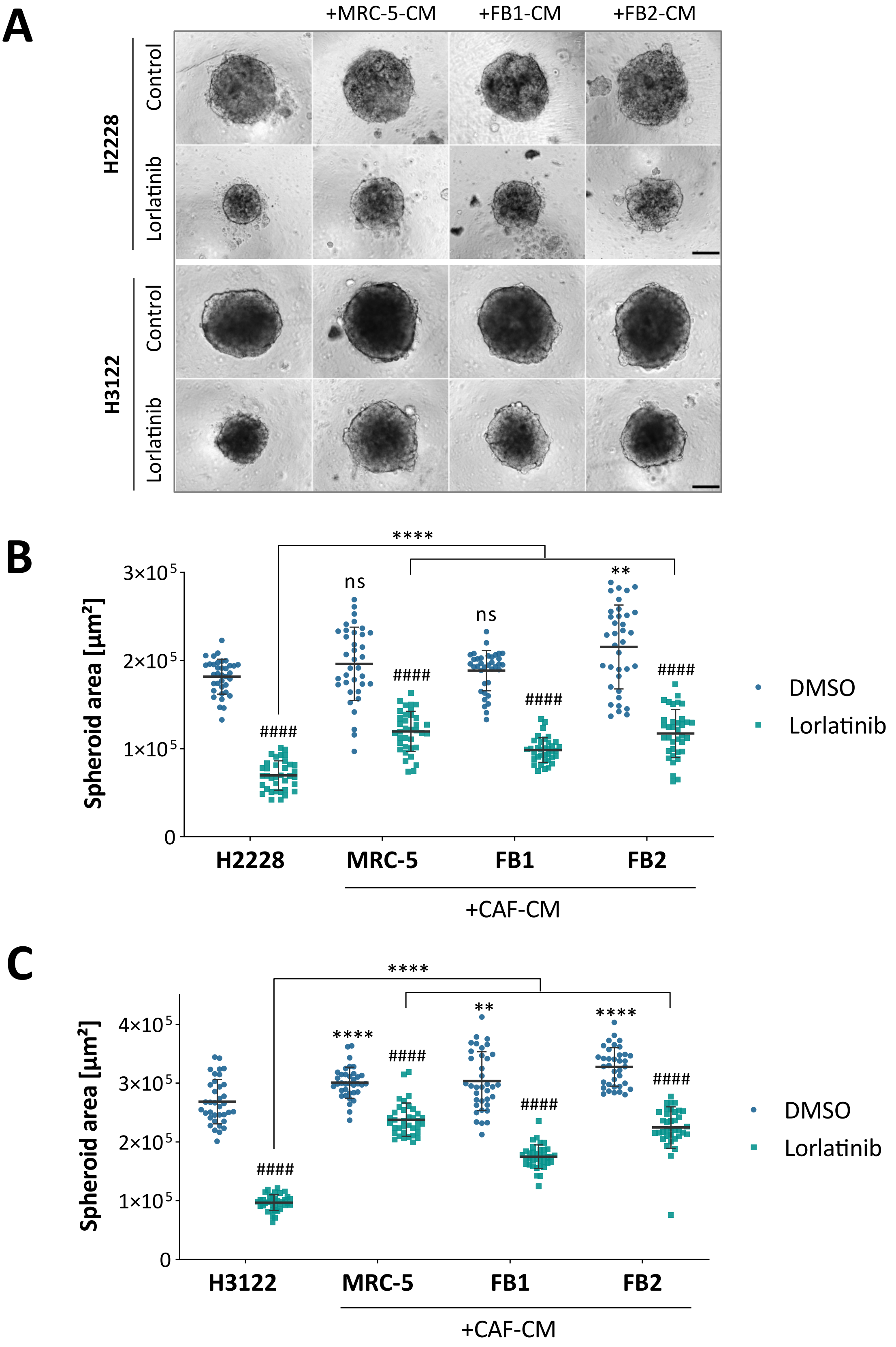


***Supplementary Figure S9:*** (A) Representative phase contrast images of homotypic H2228 and H3122 lung tumor spheroids treated according to the indicated conditions for 72 h. Scale bar: 200 µm. Changes in spheroid size following lorlatinib treatment in the absence or presence of FB2-CAF-conditioned medium were investigated by analyzing spheroid areas of imaged H2228 (B) and H3122 (C) lung tumor spheroids. Spheroid size values of twelve individual spheroids are given per experimental group (n = 3). ####, p≤0.0001 compared to DMSO controls. **, p≤0.01; ****, p≤0.0001 in comparison to lorlatinib-treatment alone.

***Supplementary Table S1:*** *Average IC50- and IC75-values of dose-response curves*

| | Brigatinib | | Lorlatinib | | |
|---|---|---|---|---|---|
| | $IC_{50}$ | $IC_{75}$ | $IC_{50}$ | $IC_{75}$ | |
| **H2228** | 15.92 | 77.77 | 3.39 | 44.65 | [nM] |
| **H3122** | 33.22 | 151.03 | 8.27 | 39.15 | [nM] |
| **A549** | 1.99 | 5.59 | 226.63 | 2395.93 | [µM] |

$IC_{50}$- and $IC_{75}$-values refer to drug concentrations that correspond to a 50% and 75% reduction in cell viability, respectively, compared to untreated control. Values are given as mean derived from two independent experiments. $IC_{50}$, half-maximal inhibitory concentration; $IC_{75}$, 75% inhibitory concentration.

***Supplementary Table S2:*** Quality control metrics following scRNA-sequencing applying the 10x Genomics plattform

| Cell line | Sample | Achieved cell recovery | Mean reads per cell | Median genes per cell | Reads mapped confidently to genome [%] | Number of cells remaining after processing |
|---|---|---|---|---|---|---|
| **H2228** | DMSO | 6,54 | 49,892 | 4,668 | 86.1 | 5,167 |
| | Brigatinib | 2,126 | 165,777 | 6,688 | 84.6 | 1,369 |
| **H2228 + FB2** | DMSO | 7,987 | 43,952 | 4,345 | 86.8 | 6,93 |
| | Brigatinib | 7,747 | 40,547 | 3,832 | 81.0 | 6,651 |
| **H3122** | DMSO | 8,343 | 41,773 | 3,296 | 86.7 | 5,867 |
| | Brigatinib | 3,355 | 102,37 | 4,698 | 82.8 | 2,5 |
| **H3122 + FB2** | DMSO | 6,205 | 54,727 | 3,268 | 88.0 | 3,764 |
| | Brigatinib | 8,203 | 40,54 | 2,677 | 83.5 | 6,17 |
| **H2228** | DMSO | 7,083 | 48,732 | 4,637 | 84.3 | 4,517 |
| | Lorlatinib | 3,135 | 112,604 | 5,476 | 85.4 | 1,989 |
| **H2228 + FB2** | DMSO | 5,386 | 63,431 | 5,153 | 85.0 | 4,656 |
| | Lorlatinib | 6,653 | 50,136 | 4,331 | 86.0 | 5,558 |
| **H3122** | DMSO | 3,951 | 78,067 | 4,781 | 86.5 | 3,067 |
| | Lorlatinib | 2,408 | 153,914 | 5,682 | 86.2 | 1,939 |
| **H3122 + FB2** | DMSO | 3,985 | 90,315 | 5,543 | 88.2 | 3,438 |
| | Lorlatinib | 3,529 | 97,475 | 5,052 | 86.5 | 2,793 |

***Supplementary Table S3:*** *Predicted interaction pairs obtained via ligand-receptor interaction analysis using 'RNA-magnet'.*

| Predicted paracrine interaction pairs | | |
|---|---|---|
| **Score** | **Ligand** | **Receptor** |
| 0.707 | HGF | SDC1 |
| 0.627 | WNT5A | LRP5 |
| 0.593 | HGF | MET |
| 0.593 | IGFBP4 | LRP6 |
| 0.580 | HGF | ST14 |
| 0.478 | WNT5A | RYK |
| 0.473 | WNT5A | LRP6 |
| 0.452 | TNFRSF11B | TNFSF10 |
| 0.435 | SFRP1 | FZD6 |
| 0.426 | FGF2 | SDC1 |
| 0.363 | WNT5A | FZD5 |
| 0.316 | WNT5A | FZD6 |
| 0.311 | NRG1 | ERBB3 |
| 0.279 | BDNF | SORT1 |
| 0.273 | FGF2 | SDC4 |

***Supplementary Table S4:*** *Oligonucleotides (primer) used for real-time PCR*

| Primer | Sequence | Application | Supplier |
|---|---|---|---|
| ACACA | forward 5'- -3' TGAACTTCACACAGGTAGTCTGCC<br>reverse 5'- -3' TGGAACACTCGATGGAGTTTCT | SYBR | Sigma-Aldrich |
| ACLY | forward 5'- -3' GAGGCATATCCAGAGGAAGCC<br>reverse 5'- -3' TCCCTTTGGGGTTCAGCAAG | SYBR | Sigma-Aldrich |
| ACTA2 | forward 5'- -3' CTGTTCCAGCCATCCTTCAT<br>reverse 5'- -3' TCATGATGCTGTTGTAGGTGGT | #58 UPL | Sigma-Aldrich |
| ACTB | forward 5'- -3' CCAACCGCGAGAAGATGA<br>reverse 5'- -3' CCAGAGGCGTACAGGGATAG | #64 UPL | Sigma-Aldrich |
| FAP | forward 5'- -3' TGGCGATGAACAATATCCTAGA<br>reverse 5'- -3' ATCCGAACAACGGGATTCTT | #19 UPL | Sigma-Aldrich |
| FASN | forward 5'- -3' TGGCGATGAACAATATCCTAGA<br>reverse 5'- -3' GCAGCTGTGACACCTTCAGG | SYBR | Sigma-Aldrich |
| GAPDH | forward 5'- -3' AGCCACATCGCTCAGACAC<br>reverse 5'- -3' GCCCAATACGACCAAATCC | #60 UPL | Sigma-Aldrich |
| SCD1 | forward 5'- -3' TAAGTTGGAGACGATGCCCC<br>reverse 5'- -3' TGGGCCTTCCTTATCCTTGT | SYBR | Sigma-Aldrich |
| SREBP-1c | forward 5'- -3' GGAGCCATGGATTGCACTTT<br>reverse 5'- -3' TCAAATAGGCCAGGGAAGTCA | SYBR | Sigma-Aldrich |
